# Unlocking a signal of introgression from codons in Lachancea kluyveri using a mutation-selection model

**DOI:** 10.1101/539148

**Authors:** Cedric Landerer, Brian C. O’Meara, Russell Zaretzki, Michael A. Gilchrist

## Abstract

For decades, codon usage has been used as a measure of adaptation for translational efficiency and translation accuracy of a gene’s coding sequence. These patterns of codon usage reflect both the selective and mutational environment in which the coding sequences evolved. Over this same period, gene transfer between lineages has become widely recognized as an important biological phenomenon. Nevertheless, most studies of codon usage implicitly assume that all genes within a genome evolved under the same selective and mutational environment, an assumption violated when introgression occurs. In order to better understand the effects of introgression on codon usage patterns and vice versa, we examine the patterns of codon usage in *Lachancea kluyveri*, a yeast which has experienced a large introgression. We quantify the effects of mutation bias and selection for translation efficiency on the codon usage pattern of the endogenous and introgressed exogenous genes using a Bayesian mixture model, ROC SEMPPR, which is built on mechanistic assumptions about protein synthesis and grounded in population genetics.

We find substantial differences in codon usage between the endogenous and exogenous genes, and show that these differences can be largely attributed to differences in mutation bias favoring A/T ending codons in the endogenous genes while favoring C/G ending codons in the exogenous genes. Recognizing the two different signatures of mutation bias and selection improves our ability to predict protein synthesis rate by 42% and allowed us to accurately assess the decaying signal of endogenous codon mutation and preferences. In addition, using our estimates of mutation bias and selection, we identify *Eremothecium gossypii* as the closest relative to the exogenous genes, providing an alternative hypothesis about the origin of the exogenous genes, estimate that the introgression occurred ∼ 6 × 10^8^ generation ago, and estimate its historic and current selection against mismatched codon usage.

Our work illustrates how mechanistic, population genetic models like ROC SEMPPR can separate the effects of mutation and selection on codon usage and provide quantitative estimates from sequence data.

## Background

Synonymous codon usage patterns varies within a genome and between taxa, reflecting differences in mutation bias, selection, and genetic drift. The signature of mutation bias is largely determined by the organism’s internal or cellular environment, such as their DNA repair genes or UV exposure. While this mutation bias is an omnipresent evolutionary force, its impact can be obscured or amplified by selection. The signature of selection on codon usage is largely determined by an organism’s cellular environment alone, such as, but not limited to, its tRNA species, their copy number, and their post-transcriptional modifications. In general, the strength of selection on codon usage is assumed to increase with its expression level (Gouy and Gautier, 1982; Ikemura, 1985; Bulmer, 1990), specifically its protein synthesis rate (Gilchrist, 2007). Thus as protein synthesis increases, codon usage shifts from a process dominated by mutation to a process dominated by selection. The overall efficacy of mutation and selection on codon usage is a function of the organism’s effective population size *N*_*e*_. ROC SEMPPR allows us to disentangle the evolutionary forces responsible for the patterns of codon usage bias (Sharp and Li, 1987; Wright, 1990; M *et al*., 1993) (CUB) encoded in an species’ genome, by explicitly modeling the combined evolutionary forces of mutation, selection, and drift (Gilchrist, 2007; Shah and Gilchrist, 2011; Wallace *et al*., 2013; Gilchrist *et al*., 2015). In turn, these evolutionary parameters should provide biologically meaningful information about the lineage’s historical cellular and external environment.

Most studies implicitly assume that the CUB of a genome is shaped by a single cellular and external environment. However, this assumption is clearly violated to increasing degrees via horizontally gene transfer, large scale introgressions, and hybrid specie formation. In these scenarios, one would expect to see the signature of multiple cellular environments in a genome’s CUB (Médigue *et al*., 1991; Lawrence and Ochman, 1997). Indeed, differences in CUB between linages have been proposed to have a major effect on their rates of gene transfer with rates declining with differences in their CUB. On a more practical level, if differences in codon usage of transferred genes are not taken into account for, they may distort the interpretation of codon usage patterns. Such distortion could lead to the wrong inference of codon preference for an amino acid (Shah and Gilchrist, 2011; Gilchrist *et al*., 2015), underestimate the variation in protein synthesis rate, or distort estimates of mutation bias when analyzing a genome.

To illustrate these ideas, we analyze the CUB of the genome of the yeast *Lachancea kluyveri* using ROC SEMPPR, a population genetics based model of synonymous codon usage evolution that accounts for and, in turn, can estimate the contribution of of mutation bias Δ*M*, selection bias. The mathematics of ROC SEMPPR are derived on a mechanistic description of ribosome movement along an mRNA, although the approximation of other biological mechanisms could also be consistent with the model. Broadly speaking, ROC SEMPPR allows us to quantify the cellular environment in which genes have evolved by separately estimating the effects of mutation bias and selection bias on codon usageDE between synonymous codons and protein synthesis rate *ϕ* to the patterns of codon usage observed within a set of genes. Briefly, the set of Δ*M* for an amino acid quantifies the relative differences in mutational stability or bias between the synonymous codons of the amino acid 𝕊. In the absence of selection bias (or equivalently when gene expression *ϕ* = 0), the equilibrium frequency of synonymous codon *i* is simply exp[−Δ*M* _*i*_]/(*∑* _*j∈*𝕊_ exp[−Δ*M* _*j*_]. Because the time units of protein production rate have no intrinsic time scale, we define the average protein production rate for a set of genes to be one, i.e. 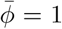 by definition (Gilchrist *et al*., 2015). In order to facilitate comparisons between gene sets, we express both, Δ*M* and Δ*η*, as deviation from the mean of each synonymous codon family (see Materials and Methods for details). Nevertheless, the difference Δ*η* describes the difference in fitness between two synonymous codons relative to drift for a gene whose protein production rate *ϕ* is equal to the the average rate of protein production 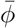 across the set of genes. In other words, for a gene whose protein is expressed at the average rate, for any two given synonymous codons *i* and *j*, Δ*η*_*i*_ − Δ*η*_*j*_ = *N*_*e*_s.

The *Lachancea* clade diverged from the Saccharomyces clade, prior to its whole genome duplication ∼100 Mya ago (Marcet-Houben and Gabaldón, 2015; Beimforde *et al*., 2014). Since that time, *L. kluyveri*, which is sister species to all other *Lachancea spp*., has experienced a large introgression of exogenous genes (1 Mb, 457 genes) which is found in all of its populations (Payen *et al*., 2009; Friedrich *et al*., 2015), but in no other known Lachancea species (Vakirlis *et al*., 2016). The introgression replaced the left arm of the C chromosome and displays a 13% higher GC content than the endogenous *L. kluyveri* genome (Payen *et al*., 2009; Friedrich *et al*., 2015). Previous studies suggest that the source of the introgression is probably a currently unknown or potentially extinct Lachancea lineage based on gene concatenation or synteny relationships (Payen *et al*., 2009; Friedrich *et al*., 2015; Vakirlis *et al*., 2016; Brion *et al*., 2017). These characteristics make *L. kluyveri* an ideal model to study the effects of an introgressed cellular environment and the resulting mismatch in codon usage.

While previous studies (Shah and Gilchrist, 2011; Wallace *et al*., 2013) have used information on gene expression to separate the effects of mutation and selection on codon usage, ROC SEMPPR does not need such information but can provide it. ROC SEMPPR’s resulting predictions of protein synthesis rates have been shown to be on par with laboratory measurements (Shah and Gilchrist, 2011; Gilchrist *et al*., 2015). In contrast to often used heuristic approaches to study codon usage (Sharp and Li, 1987; Wright, 1990; dos Reis *et al*., 2004), ROC SEMPPR explicitly incorporates and distinguishes between mutation and selection effects on codon usage and properly weights its estimates by amino acid usage (Cope *et al*., 2018). We use ROC SEMPPR to separately describe the two cellular environments reflected in the *L. kluyveri* genome; the signature of the endogenous environment reflected in the larger set of non-introgressed genes and the decaying signature of the ancestral, exogenous environment in the smaller set of introgressed genes. Our results indicate that the current difference in GC content between endogenous and exogenous genes is mostly due to the differences in mutation bias Δ*M* of their respective cellular environments. Taking the different signatures of Δ*M* and selection bias Δ*η* of the endogenous and exogenous sets of genes substantially improves our ability to predict present day protein synthesis rates *ϕ*. These endogenous and exogenous gene set specific estimates of Δ*M* and Δ*η*, in turn, allow us to address more refined biological questions. For example, we find support for an alternative origin of the exogenous genes and identify *E. gossypii* as the nearest sampled relative of the source of the introgressed genes out of the 332 budding yeast lineages with sequenced genomes (Shen *et al*., 2018). While this inference is in contrast to previous work (Payen *et al*., 2009; Friedrich *et al*., 2015; Vakirlis *et al*., 2016; Brion *et al*., 2017), we find additional phylogenetic support for via gene tree reconstruction and gene synteny. We also estimate the age of the introgression to be on the order of 0.2 - 1.7 Mya, estimate the selection against these genes, both at the time of introgression and now, and predict a detectable signature of CUB to persist in the introgressed genes for another 0.3 - 2.8 Mya, highlighting the sensitivity of our approach.

## Results

### The Signatures of two Cellular Environments within *L. kluyveri* ‘s Genome

We used our software package AnaCoDa (Landerer *et al*., 2018) to compare model fits of ROC SEMPPR to the entire *L. kluyveri* genome and its genome partitioned into two sets of 4,864 endogenous and 497 exogenous genes. These two set where initially identified based on their striking difference in GC content Payen *et al*. (2009), with very little overlap in GC content between the two sets (Figure S1a). ROC SEMPPR is a statistical model that relates the effects of mutation bias Δ*M*, selection bias Δ*η* between synonymous codons and protein synthesis rate *ϕ*, to explain the observed codon usage patterns. Thus, the probability of observing a synonymous codon is proportional to *p ∝* exp(−Δ*M* −Δ*ηϕ*) (Gilchrist *et al*., 2015). Briefly, Δ*M* describes the mutation bias between two synonymous codons at stationarity under a time reversible mutation model. Because ROC SEMPPR only considers the stationary probabilities, only variation in mutation bias, not absolute mutation rates can be detected. Δ*η* describes the fitness difference between two synonymous codons relative to drift (Gilchrist *et al*., 2015). Since Δ*η* is scaled by protein synthesis rate *ϕ*, this term is dominant in highly expressed genes and tends towards 0 in low expression genes, allowing us to separate the effect of mutation bias and selection bias on codon usage. We express both, Δ*M* and Δ*η*, as deviation from the mean of each synonymous codon family which prevents that the choice of the reference codon affects our results (see Materials and Methods for details).

Bayes factor strongly support the hypothesis that the *L. kluyveri* genome consists of genes with two different and distinct patterns of codon usage bias rather than a single (*K* = exp(42, 294); Table 1). We find additional support for this hypothesis when we compare our predictions of protein synthesis rate to empirically observed mRNA expression values as a proxy for protein synthesis. Specifically, we improve the variance explained by our predicted protein synthesis rates by ∼42%, from *R*^2^ = 0.33 (*p* < 10^10^) to 0.46 (*p* < 10^10^) (Figure 1). While the implicit consideration of GC content in this analysis certainly plays a roll, it does not explain the improvement in *R*^2^ (Figure S1b).

**Table 1:**
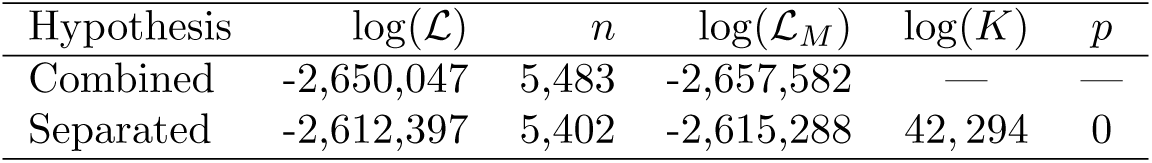
Model selection of the two competing hypothesis. Combined: mutation bias and selection bias for synonymous codons is shared between endogenous and exogenous genes. Separated: mutation bias and selection bias for synonymous codons is allowed to vary between endogenous and exogenous genes. Reported are the log-likelihood, log(*ℒ*), the number of parameters estimated *n*, the log-marginal likelihood log(ℒ_*M*_), Bayes Factor K, and the p-value of the likelihood ratio test.

**Figure 1:**
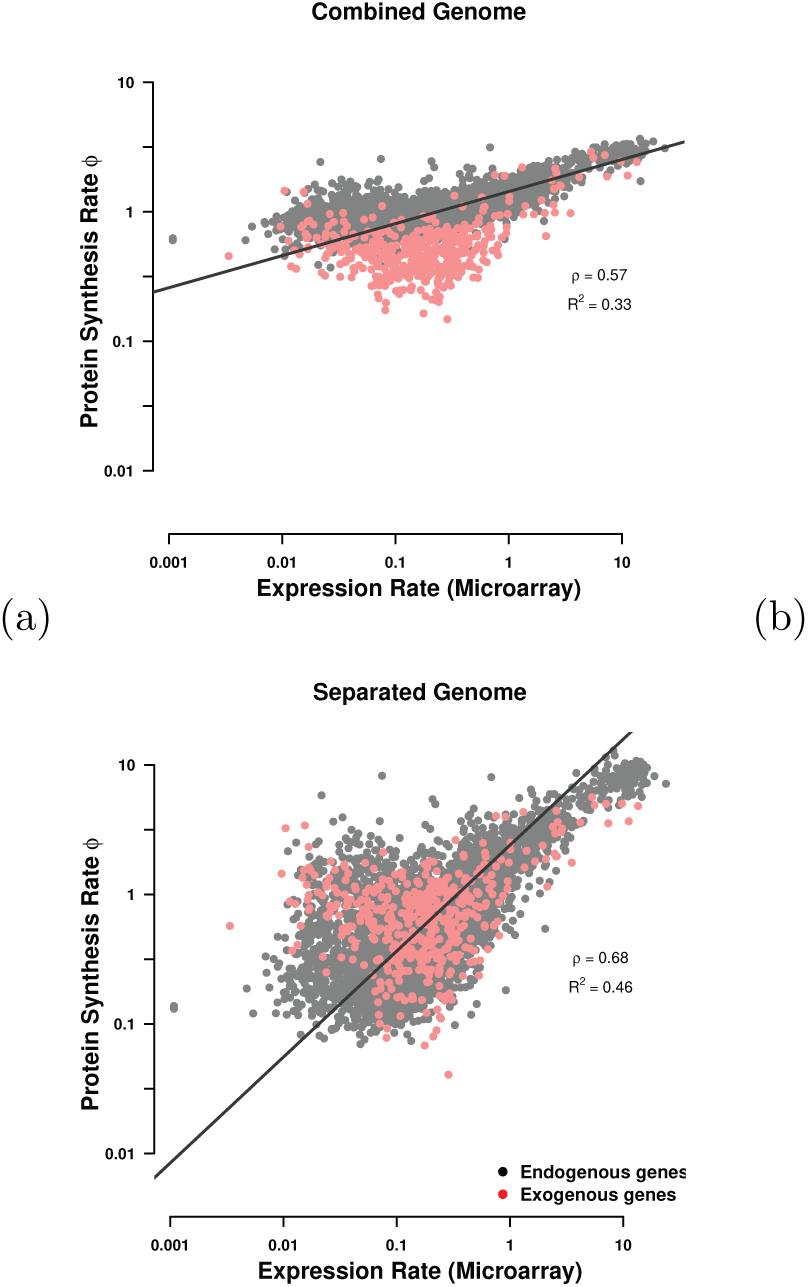
Comparison of predicted protein synthesis rate *ϕ* to mRNA abundance from Tsankov *et al*. (2010) for (a) the combined genome where mutation bias and selection bias parameters Δ*M* and Δ*η* are estimated for the combined endogenous and exogenous gene sets, and (b) where Δ*M* and Δ*η* are estimated separately for the endogenous and exogenous gene sets. Endogenous genes are displayed in black and exogenous genes in red. Black line indicates type II regression line (Sokal and Rohlf, 1981).

### Comparing Differences in the Endogenous and Exogenous Codon Usage

Because ROC SEMPPR defines 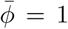, it makes the interpretation of Δ*η* as selection on codon usage of the average gene with *ϕ* = 1 straightforward and gives us the ability to compare the efficacy of selection *sN*_*e*_ across genomes. While it may be expected for the endogenous and exogenous genes to differ in the their codon usage pattern due to the large difference in GC content it is not clear how much of this difference is due to differences in the mutation bias Δ*M* or selection bias Δ*η* between the gene sets. To better understand the differences in the endogenous and exogenous cellular environments, we compared our parameter estimates of Δ*M* and Δ*η* for the two sets of genes. Our estimates of Δ*M* for the endogenous and exogenous genes were negatively correlated (*ρ* = −0.49, *p* = 3.56 × 10^−5^), indicating weak similarity with only ∼5% of the codons share the same sign between the two mutation environments (Figure 2a). Overall, the endogenous genes only show a selection preference for C and G ending codons in ∼58% of the codon families. In contrast, the exogenous genes display a strong preference for A and T ending codons in ∼89% of the codon families.

**Figure 2:**
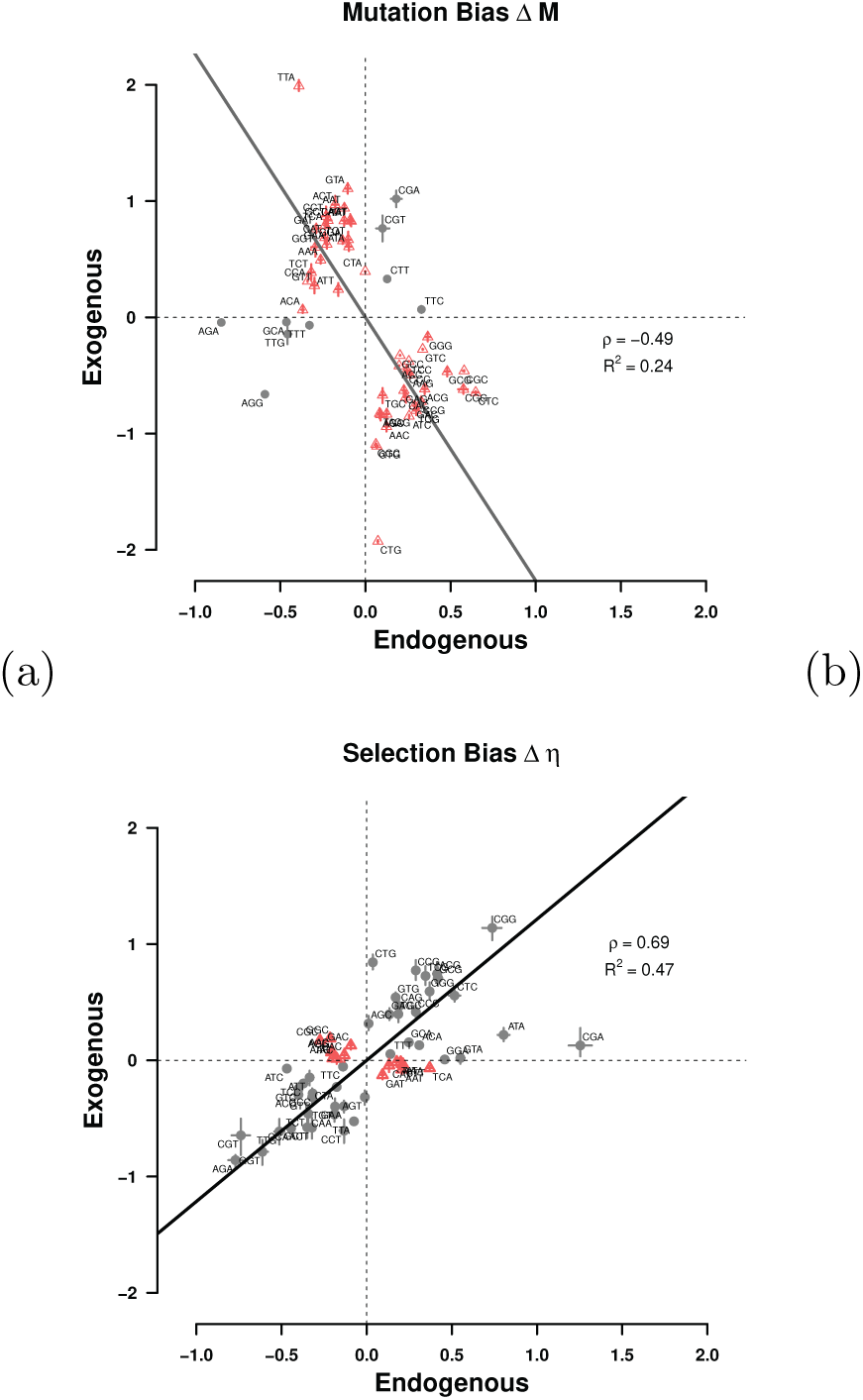
Comparison of (a) mutation bias Δ*M* and (b) selection bias Δ*η* parameters for endogenous and exogenous genes. Estimates are relative to the mean for each codon family. Black dots indicate Δ*M* or Δ*η* parameters with the same sign for the endogenous and exogenous genes, red dots indicate parameters with different signs. Black line indicates type II regression line (Sokal and Rohlf, 1981). Dashed lines mark quadrants.

For example, the endogenous genes show a mutational bias for A and T ending codons in ∼95% of the codon families The exception is Leucine (Leu, L), where mutation appears to favor the codon TTG over TTA (Figure 3, Table S1). The exogenous genes display an equally consistent mutational bias towards C and G ending codons (the exception being Phe, F). In contrast to Δ*M*, our estimates of Δ*η* for the endogenous and exogenous genes were positively correlated (*ρ* = 0.69) and showing the same sign in ∼53% of codons between the two selection environments (Figure 2).

**Figure 3:**
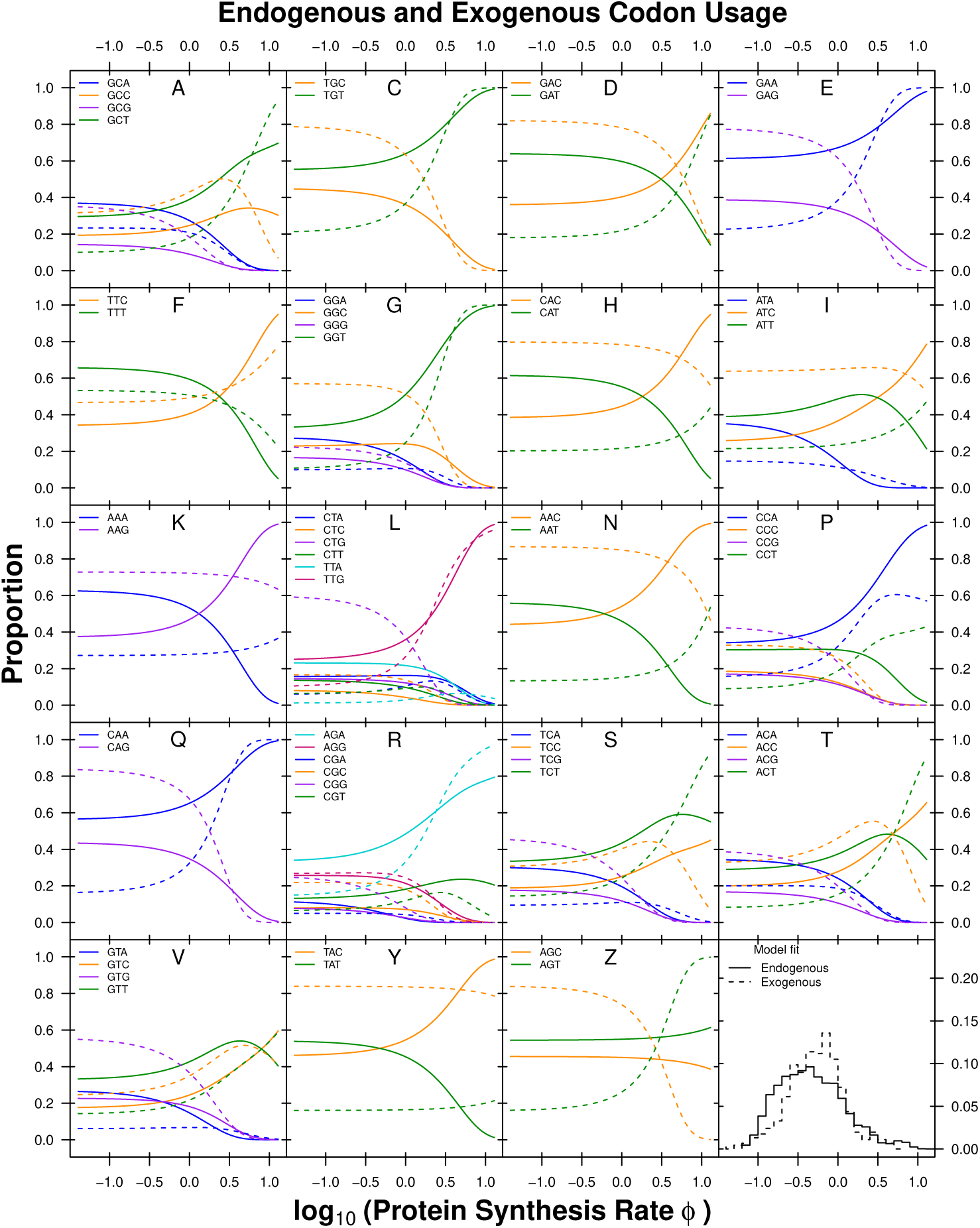
Codon usage patterns for 19 amino acids. Amino acids are indicated as one letter code. The amino acids Serine was split into two groups (S and Z) as Serine is coded for by two groups of codons that are separated by more than one mutation. Solid line indicates the endogenous codon usage, dashed line indicates the exogenous codon usage.

We find that the signature of selection bias Δ*η* also differs substantially between the endogenous and exogenous gene sets. The difference in codon usage between endogenous and exogenous genes is striking as the sign for Δ*η* changes, indicating a change in codon preference for some amino acids. As a result, our estimates of the optimal codon differ in nine cases between endogenous and exogenous genes (Figure 3, Table S2). For example, the usage of the Asparagine (Asn, N) codon AAC is increased in highly expressed endogenous genes but the same codon is depleted in highly expressed exogenous genes. For Aspartic acid (Asp, D), the combined genome shows the same codon preference in highly expressed genes as the exogenous gene set. Generally, fits to the complete *L. kluyveri* genome reveal that the relatively small exogenous gene set (∼10% of genes) has a disproportionate effect on the model fit (Figure S2, S3).

Of the nine cases in which the endogenous and exogenous genes show differences in the selectively most favored codon five cases (Asp, D; His, H; Lys, K; Asn, N; and Pro, P) the endogenous genes favor the codon with the most abundant tRNA. For the remaining four cases (Ile, I; Ser, S; Thr, T; and Val, V), there are no tRNA genes for the wobble free cognate codon encoded in the *L. kluyveri* genome. However, the codon preference of these four amino acids in the exogenous genes matches the most abundant tRNA encoded in the *L. kluyveri* genome. In contrast to Δ*M*, our estimates of selection bias Δ*η* for the endogenous and exogenous genes are positively correlated (*ρ* = 0.69, *p* = 9.76 × 10^−10^) and show the same sign in ∼53% of the cases (Figure 2).

This striking difference in codon usage was noted previously. For example, using RSCU (Sharp and Li, 1987), GAA (coding for Glu, E) was identified as the optimal synonymous codon in the whole genome and GAG as the optimal codon in the exogenous genes Payen *et al*. (2009). Our results, however, indicate that GAA is the optimal codon in both, endogenous and exogenous genes, and that the high RSCU in the exogenous genes of GAG is driven by mutation bias (Table S1 and S2). Similar effects are observed for other amino acids.

The effect of the small exogenous gene set on the fit to the complete *L. kluyveri* genome is smaller for our estimates of selection bias Δ*η* than Δ*M*, but still large. We find that the complete *L. kluyveri* genome is estimated to share the selectively preferred codon with the exogenous genes in ∼60% of codon families that show dissimilarity between endogenous and exogenous genes. We also find that the complete *L. kluyveri* genome fit shares mutationally preferred codons with the exogenous genes in ∼78% of the 19 codon families showing a difference in mutational codon preference between the endogenous and exogenous genes. In two cases, Isoleucine (Ile, I) and Arginine (Arg, R), the strong dissimilarity in mutation preference results in an estimated codon preference in the complete *L. kluyveri* genome that differs from both the endogenous, and the exogenous genes. These results clearly show that it is important to recognize the difference in endogenous and exogenous genes and treat these genes as separate sets to avoid the inference of incorrect synonymous codon preferences and better predict protein synthesis.

### Can Codon Usage Help Determine the Source of the Exogenous Genes

Since the origin of the exogenous genes is currently unknown, we explored if the information on codon usage extracted from the exogenous genes can be used to identify a potential source lineage. We combined our estimates of mutation bias Δ*M* and selection bias Δ*η* with synteny information and searched for potential source lineages of the introgressed exogenous region. We used Δ*M* to identify candidate lineages as the endogenous and exogenous genes show greater dissimilarity in mutation bias than in selection bias. We examined 332 budding yeasts (Shen *et al*., 2018) and, identified the ten lineages with the highest correlation to the exogenous Δ*M* parameters as potential source lineages (Figure 4, Table 2). Two of the ten candidate lineages utilize the alternative yeast nuclear code (NCBI codon table 12). In this case, the codon CTG codes for Serine instead of Leucine. We therefore excluded the Leucine codon family from our comparison of codon families; however, there was no need to exclude Serine as CTG is not a one step neighbor of the remaining Serine codons. A mutation between CTG and the remaining Serine codons would require two mutations with one of them being non-synonymous, which would violate the weak mutation assumption of ROC SEMPPR.

**Table 2:**
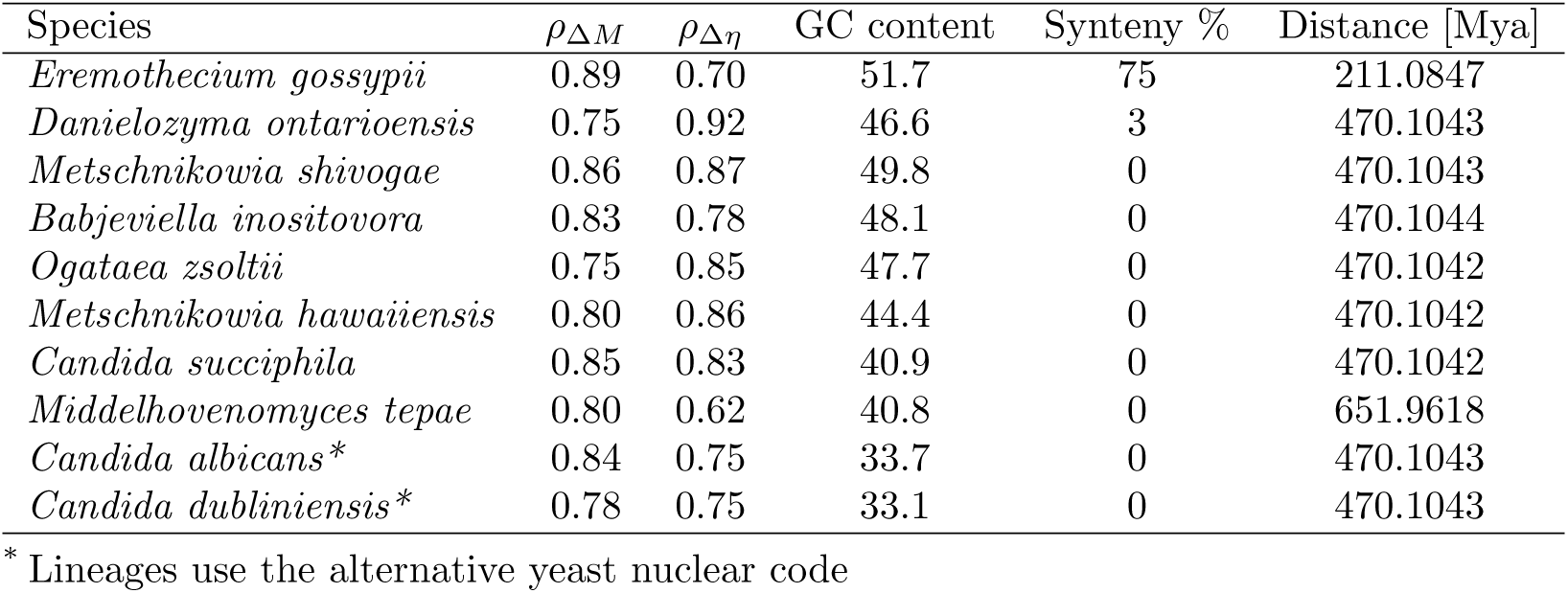
Budding yeast lineages showing similarity in codon usage with the exogenous genes. *ρ*_Δ*M*_ and *ρ*_Δ*η*_ represent the Pearson correlation coefficient for exogenous Δ*M* and Δ*η* with the indicated species’, respectively. GC content is the average GC content of the whole genome. Synteny is the percentage of the exogenous genes found in the listed lineage. Only one lineage (*E. gossypii*) shows a similar GC content > 50%.

**Figure 4:**
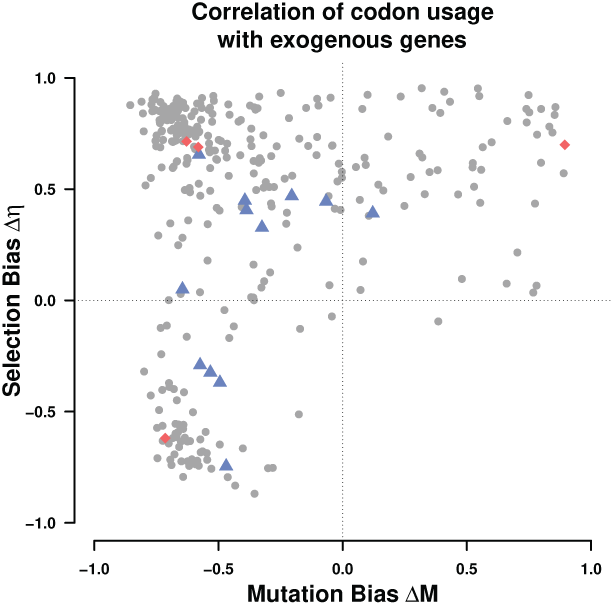
Correlation coefficients of Δ*M* and Δ*η* of the exogenous genes with 332 examined budding yeast lineages. Dots indicate the correlation of Δ*M* and Δ*η* of the lineages with the exogenous parameter estimates. Blue triangles indicate the *Lachancea* and red diamonds indicate *Eremothecium* species. All regressions were performed using a type II regression (Sokal and Rohlf, 1981).

The endogenous *L. kluyveri* genome exhibits codon usage very similar to most (77 %) yeast lineages examined, indicating that most of the examined yeasts share a similar codon usage (Figure S4). Only ∼17% of all examined yeast show a positive correlation in both, Δ*M* and Δ*η* with the exogenous genes, whereas the vast majority of lineages (∼83%) show a negative correlation for Δ*M*, only 21 % show a negative correlation for Δ*η*.

Comparing synteny between the exogenous genes, which are restricted to the left arm of chromosome C, and the candidate yeast species we find that *E. gossypii* is the only species that displays high synteny (Table 2). Furthermore, the synteny relationship between the exogenous region and other yeasts appears to be limited to Saccharomycetaceae clade. Given these results, we conclude that, of the 332 examined yeast lineages the *E. gossypii* lineage is the most likely source of the introgressed exogenous genes. Previous studies which studied the exogenous genes and chromosome recombination in the Lachancea clade concluded that the exogenous region originated from within the Lachancea clade, from an unknown or potentially extinct lineage (Payen *et al*., 2009; Friedrich *et al*., 2015; Vakirlis *et al*., 2016). While it is not possible for us to dispute this hypothesis, our results provide a novel hypothesis about the origin of the exogenous genes.

To further test the plausibility of *E. gossypii* as potential source linage, we identified 127 genes in our dataset (Shen *et al*., 2018) with homologous genes in *E. gossypii* and other Lachancea and used IQTree (Nguyen *et al*., 2015) to infer the phylogenetic relationship of the exogenous genes. Our results show that at least ∼45% of exogenous genes (57/127) are more closely related to *E. gossypii* than to other Lachancea S5. Interestingly, our results also indicate that codon usage does not necessarily correlate with phylogenetic distance (Table 2).

### Estimating Introgression Age

If we assume that the exogenous genes originated from the *E. gossypii* lineage, we can estimate the age of the introgression based on our estimates of mutation bias Δ*M*. We modeled the change in codon frequency over time as exponential decay, and estimated the age of the introgression assuming that *E. gossypii* still represents the mutation bias of its ancestral source lineage at the time of the introgression and a constant mutation rate. We infer the age of the introgression to be on the order of 6.2 ± 1.2 × 10^8^ generations. Assuming *L. kluyveri* experiences between one and eight generations per day, we estimate the introgression to have occurred between 212, 000 to 1, 700, 000 years ago. Our estimate places the time of the introgression earlier than the previous estimate of 19,000 - 150,000 years by Friedrich *et al*. (2015).

Using our model of exponential decay model, we also estimated the persistence of the signal of the exogenous cellular environment. We predict that the Δ*M* signal of the source cellular environment will have decayed to be within one percent of the *L. kluyveri* environment in ∼5.4 ± 0.2 × 10^9^ generations, or between 1, 800, 000 and 15, 000, 000 years. Together, these results indicate that the mutation signature of the exogenous genes will persist for a very long time.

### Estimating Selection against Codon Mismatch of the Exogenous Genes

We define the selection against inefficient codon usage as the difference between the fitness on the log scale of an expected, replaced endogenous gene and the exogenous gene, *s ∝ ϕ*Δ*η* due to the mismatch in codon usage parameters (See Methods for details). As the introgression occurred before the diversification of *L. kluyveri* and has fixed throughout all populations (Friedrich *et al*., 2015), we can not observe the original endogenous sequences that have been replaced by the introgression. Overall, we predict that a small number of low expression genes (*ϕ* < 1) were weakly exapted at the time of the introgression (Figure 5a). Thus, they appear to provide a small fitness advantage due to the accordance of exogenous mutation bias with endogenous selection bias (compare Figure S2 and S3). High expression genes (*ϕ* > 1) are predicted to have faced the largest selection against their mismatched codon usage in the novel cellular environment. In order to account for differences in the efficacy of selection on codon usage either due to the cost of pausing, differences in the effective population size, or the decline in fitness with every ATP wasted between the donor lineage and *L. kluyveri* we added a linear scaling factor *κ* to scale our estimates of Δ*η* between the donor lineage and *L. kluyveri* and searched for the value that minimized the cost of the introgression, thus giving us the best case scenario (See Methods for details).

**Figure 5:**
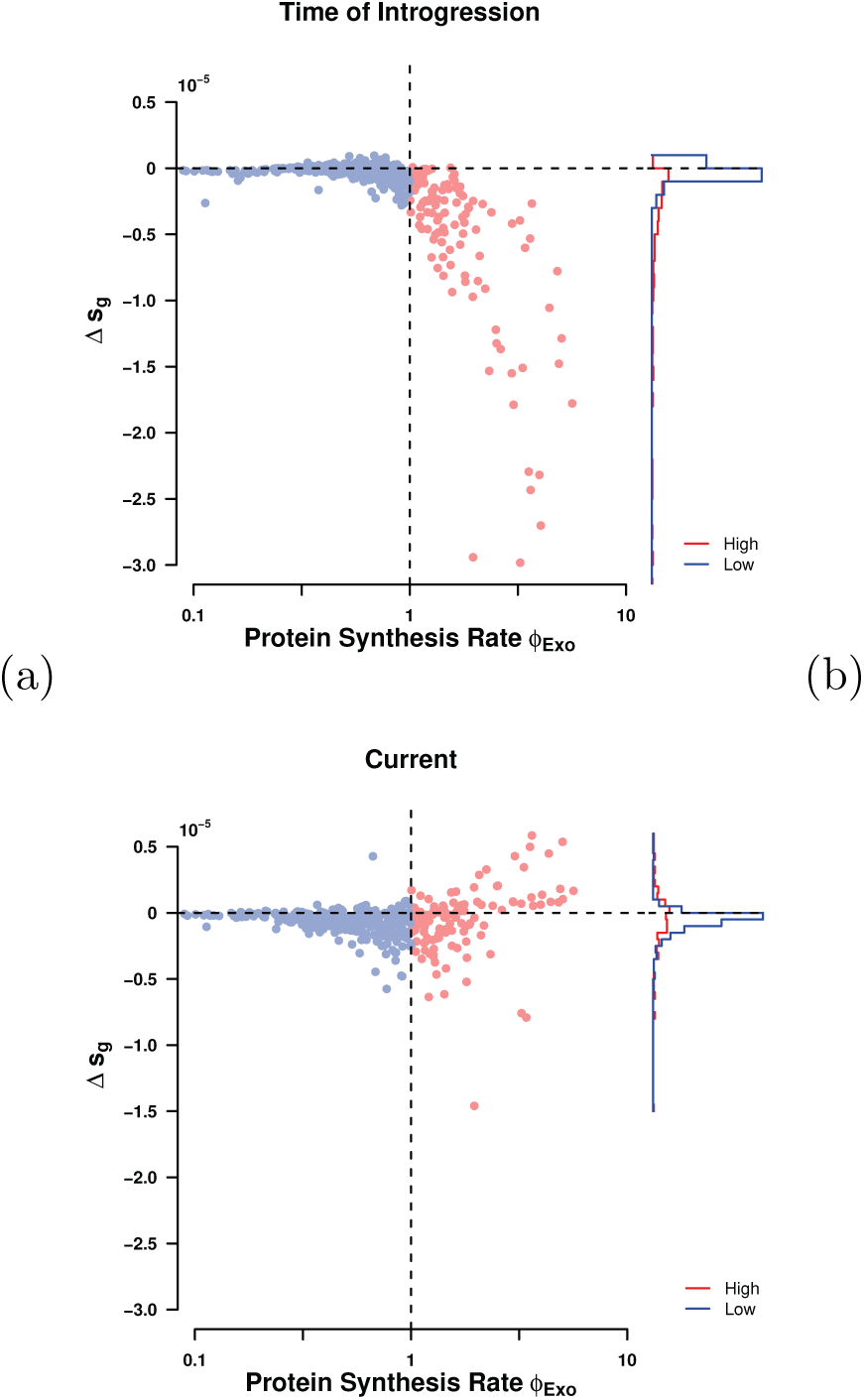
Selection against mismatched codon usage *s* = Δ*ηϕ* (a) at the time of introgression (*κ* = 5), and (b) currently (*κ* = 1). Vertical dashed line indicates split between high and low expression genes at *ϕ* = 1. Horizontal dashed line indicates neutrality.

Using our estimates of Δ*M* and Δ*η* from the endogenous genes and assuming the current exogenous amino acid composition of genes is representative of the replaced endogenous genes, we estimate the strength of selection against the exogenous genes at the time of introgression (Figure 5a) and currently (Figure 5b). Estimates of selection bias for the exogenous genes show that, while well correlated with the endogenous genes, only nine amino acids share the same selectively preferred codon. Exogenous genes are, therefore, expected to represent a significant reduction in fitness for *L. kluyveri* due to mismatch in codon usage. Since Δ*η* is proportional to the difference in fitness between the wild type and a mutant, we can use our estimates of Δ*η* to approximate the selection against the exogenous genes Δ*s* (Sella and Hirsh, 2005; Gilchrist *et al*., 2015). We estimate that the selection against all exogenous genes due to mismatched codon usage to have been Δ*s* ≈− 0.0008 at the time of the introgression and ≈−0.0003 today. This reduction in Δ*s* is primarily due to adaptive changes to the codon usage of the most highly expressed, introgressed genes (Figures 5a & S8). Based on the selection against the codon mismatch at the time of the introgression and assuming an effective population size *N*_*e*_ on the order of 10^7^ (Wagner, 2005), we estimate a fixation probability of (1 − exp[−Δ*s*])/(1 − exp[−2Δ*sN*_*e*_]) ≈ 10^−6952^ (Sella and Hirsh, 2005) for the exogenous genes. Clearly, the possibility of fixation under this simple scenario is effectively zero. In order for the exogenous genes to have reached fixation one or more exogenous loci must have provided a selective advantage not considered in this study (See Discussion).

## Discussion

In order to study the evolutionary effects of the large scale introgression of the left arm of chromosome C, we used ROC SEMPPR, a mechanistic model of ribosome movement along an mRNA. The usage of a mechanistic model rooted in population genetics allows us generate more nuanced quantitative parameter estimates and separate the effects of mutation and selection on the evolution of codon usage. This allowed us to calculate the selection against the introgression, and provides *E. gossypii* as a potential source lineage of the introgression which was previously not considered. Our parameter estimates indicate that the *L. kluyveri* genome contains distinct signatures of mutation and selection bias from both an endogenous and exogenous cellular environment. By fitting ROC SEMPPR separately to *L. kluyveri* ‘s endogenous and exogenous sets of genes we generate a quantitative description of their signatures of mutation bias and natural selection for efficient protein translation.

In contrast to other methods such as RSCU, CAI, or tAI, ROC SEMPPR does not rely on external information such as gene expression or tRNA gene copy number Sharp and Li (1987); dos Reis *et al*. (2004). Instead, ROC SEMPPR allows for the estimation of protein synthesis rate *ϕ* and separates the effects of mutation and selection on codon usage. In addition, Cope *et al*. (2018) showed that approaches like CAI are sensitive to amino acid composition, another property that distinguishes the endogenous and exogenous genes Payen *et al*. (2009).

Previous work by Payen *et al*. (2009) showed an increased bias towards GC rich codons in the exogenous genes but our results provide more nuanced insights by separating the effects of mutation bias and selection. We are able to show that the difference in GC content between endogenous and exogenous genes is mostly due to differences in mutation bias as 95% of exogenous codon families show a strong mutation bias towards GC ending codons (Table S1). However, the exogenous genes show a selective preference for AT ending codons for 90% of codon families (Table S2). Acknowledging the increased mutation bias towards GC ending codons and the difference in strength of selection between endogenous and exogenous genes by separating them also improves our estimates of protein synthesis rate *ϕ* by 42% relative to the full genome estimate (*R*^2^ = 0.46, *p* = 0 vs. 0.32, *p* = 0, respectively).

Previous studies showed that nucleotide composition can be strongly affected by biased gene conversion, which, in turn would affect codon usage. Biased gene conversion is thought to act similar to directional selection, typically favoring the fixation of G/C alleles Nagylaki (1983a,b). Further, (Harrison and Charlesworth, 2011, Harrison & Charlesworth) suggested that biased gene conversion affects codon usage in S. cerevisiae. ROC SEMPPR, however, does not explicitly account for biased gene conversion. If biased gene conversion is independent of gene expression, as in the case of DNA repair, it will be absorbed in our estimates of Δ*M*. If instead biased gene conversion forms hotspots, and thus becomes gene specific, it will affect our estimates of protein synthesis *ϕ*. This might be the case at recombination hotspots. Recombination, however, is very low in the introgressed region (discussed below) Payen *et al*. (2009); Brion *et al*. (2017). The low recombination rate also indicates that the GC content had to be high before the introgression occurred.

The estimates of mutation and selection bias parameters, Δ*M* and Δ*η*, are obtained under an equilibrium assumption. Given that the introgression is still adapting to its new environment, this assumption is clearly violated. However, the adaptation of the exogenous genes progresses very slowly as a quasi-static process as shown in this work as well as Friedrich *et al*. (2015). Therefore, the genome can be assumed to maintain an internal equilibrium at any given time. We see empirical evidence for this behavior in our ability to predict gene expression and to correctly identify the low expression genes (Figure 1b).

Despite the violation of the equilibrium assumption, the mutation and selection bias parameters Δ*M* and Δ*η* of the introgressed exogenous genes contain information, albeit decaying, about its previous cellular environment. We selected the top ten lineages with the highest similarity in Δ*M* to see if our parameters estimates would allow us to identify a potential source lineage. The synteny relationship of these lineages with the exogenous genes was calculated as a point of comparison as it provides orthogonal information to our parameter estimates. Synteny with the exogenous genes is limited to the Saccharomycetaceae clade, excluding all of the potential source lineages identified using codon usage but *E. gossypii* (Table 2). Interestingly, this also showed that similarity in codon usage does not correlate with phylogenetic distance.

Previous work indicated that the donor lineage of the exogenous genes has to be a, potentially unknown, Lachancea lineage (Payen *et al*., 2009; Friedrich *et al*., 2015; Vakirlis *et al*., 2016; Brion *et al*., 2017). These previous results, however, are based on species rather than gene trees, ignoring the differential adaptation rate to their novel cellular environment between genes or do not consider lineages outside of the Lachancea clade. Considering the similarity in selection bias (Figure 2b) and our calculation of selection on the exogenous genes (Figure 5b), both of which are free of any assumption about the origin of the exogenous genes, a species tree estimated from the exogenous genes will be biased towards the Lachancea clade. Estimating individual gene trees rather than relying on a species tree provided further evidence that the exogenous genes could originate from a lineage that does not belong to the Lachancea clade. As we highlighted in this study, relatively small sets of genes with a signal of a foreign cellular environment can significantly bias the outcome of a study. The same holds true for phylogenetic inferences (Salichos and Rokas, 2013), and as we showed the signal of the original endogenous cellular environment that shaped CUB is at different stages of decay in high and low expression genes (Figure S8). In summary, our work does not dispute an unknown Lachancea as possible origin, but provides an alternative hypothesis based on the codon usage of the exogenous genes, phylogenetic analysis, and synteny.

In terms of understanding the spread of the introgression, we calculated the expected selective cost of codon mismatch between the *L. kluyveri* and *E. gossypii* lineages. Under our working hypothesis, the majority of the introgressed would have imposed a selective cost due to codon mismatch. Nevertheless, ∼30% of low expression exogenous genes (*ϕ* < 1) appeared to be exapted at the time of the introgression. This exaptation is due to the mutation bias in the endogenous genes matching the selection bias in the exogenous genes for GC ending codons. Our estimate of the selective cost of codon mismatch on the order of −0.0008. While this selective cost may not seem very large, assuming *L. kluyveri* had a large *N*_*e*_, the fixation probability of the introgression is the astronomically small value of ≈10^−6952^ ≈ 0. While this estimate heavily depends on the working hypothesis that the exogenous genes originated from the *E. gossypii* lineage, we can also calculate the hypothetical fixation probability if the current exogenous genes would introgres into *L. kluyveri*. Our estimate of the current selective cost of the mismatch of codon usage is on the order of −0.0003. The fixation probability of the current exogenous genes would still be astronomically small ≈10^−2609^ ≈ 0 These results are in accordance with previous work, highlighting the necessity of codon usage compatibility between endogenous and transferred exogenous genes Medrano-Soto *et al*. (2004); Tuller *et al*. (2011). Thus, the basic scenario of an introgression between two yeast species with large *N*_*e*_ and where the introgression solely imposes a selective cost due to codon mismatch is clearly too simplistic.

One or more loci with a combined selective advantage on the order of 0.0008 or greater would have made the introgression change from disadvantageous to effectively neutral or advantageous. While this scenario seems plausible, it raises the question as to why recombination events did not limit the introgression to only the adaptive loci. A potential answer is the low recombination rate between the endogenous and exogenous regions Payen *et al*. (2009); Brion *et al*. (2017). Estimates of the recombination rate as measured by crossovers (COs) for *L. kluyveri* are almost four times lower than for *S. cerevisae* and about half that of *Schizosaccharomyces pombe* (≈1.6 COs/Mb/meiosis, ≈6 COs/Mb/meiosis, ≈3 COs/Mb/meiosis) with no observed crossovers in the introgressed region Brion *et al*. (2017), and no observed transposable elements Payen *et al*. (2009). This is presumably due to the dissimilarity in GC content and/or a lower than average sequence homology between the exogenous region and the one it replaced. A population bottleneck reducing the *N*_*e*_ of the *L. kluyveri* lineage around the time of the introgression could also help explain the spread of the introgression. Compatible with these explanation is the possibility of several advantageous loci distributed across the exogenous region drove a rapid selective sweep and/or the population through a bottleneck speciation process.

Assuming *E. gossypii* as potential source lineage of the exogenous region, we illustrated how information on codon usage can be used to infer the time since the introgression occurred using our estimates of mutation bias Δ*M*. The Δ*M* estimates are well suited for this task as they are free of the influence of selection and unbiased by *N*_*e*_ and other scaling terms, which is in contrast to our estimates of Δ*η* (Gilchrist *et al*., 2015). Our estimated age of the introgression of 6.2 ± 1.2 × 10^8^ generations is ∼10 times longer than a previous minimum estimate by Friedrich *et al*. (2015) of 5.6 × 10^7^ generations, which was based on the effective population recombination rate and the population mutation parameter (Ruderfer *et al*., 2006). Furthermore, these estimates assume that the current *E. gossypii* and *L. kluyveri* cellular environment reflect their ancestral states at the time of the introgression. Thus, if the ancestral mutation environments were more similar (dissimilar) at the time of the introgression then our result is an overestimate (underestimate).

Further, the presented work provides a template to explore the evolution of codon usage. This applies not only to species who experienced an introgression but is more generally applicable to any species.

## Conclusion

Overall, our results show the usefulness of the separation of mutation bias and selection bias and the importance of recognizing the presence of multiple cellular environments in the study of codon usage. We also illustrate how a mechanistic model like ROC SEMPPR and the quantitative estimates it provides can be used for more sophisticated hypothesis testing in the future. In contrast to other approaches used to study codon usage like CAI (Sharp and Li, 1987) or tAI (dos Reis *et al*., 2004), ROC SEMPPR incorporates the effects of mutation bias and amino acid composition explicitly (Cope *et al*., 2018). We highlight potential issues when estimating codon preferences, as estimates can be biased by the signature of a second, historical cellular environment. In addition, we show how quantitative estimates of mutation bias and selection relative to drift can be obtained from codon data and used to infer the fitness cost of an introgression as well as its history and potential future.

## Abbreviations

AIC: Akaike information criterion;
CAI: Codon adaptation index;
CUB: Codon usage bias;
ROC SEMPPR: ribosome overhead costs Stochastic Evolutionary Model of Protein Production Rate;
RSCU: Relative synonymous codon usage;
tAI: tRNA adaptation index

## Methods

### Separating Endogenous and Exogenous Genes

A GC-rich region was identified by Payen *et al*. (2009) in the *L. kluyveri* genome extending from position 1 to 989,693 of chromosome C. This region was later identified as an introgression by Friedrich *et al*. (2015). We obtained the *L. kluyveri* genome from SGD Project http://www.yeastgenome.org/download-data/ (on 09-27-2014) and the annotation for *L. kluyveri* NRRL Y-12651 (assembly ASM14922v1) from NCBI (on 12-09-2014). We assigned 457 genes located on chromosome C with a location within the ∼1 Mb window to the exogenous gene set. All other 4864 genes of the *L. kluyveri* genome were assigned to the exogenous genes.

### Model Fitting with ROC SEMPPR

ROC SEMPPR was fitted to each genome using AnaCoDa (0.1.1) (Landerer *et al*., 2018) and R (3.4.1) (R Core Team, 2013). ROC SEMPPR was run from 10 different starting values for at least 250,000 iterations and thinned to keep every 50th iteration. After manual inspection to verify that the MCMC had converged, parameter posterior means, log posterior probability and log likelihood were estimated from the last 500 samples (last 10% of samples).

### Model selection

The marginal likelihood of the combined and separated model fits was calculated using a generalized harmonic mean estimator (Gronau *et al*., 2017). A variance scaling of 1.1 was used to scale the important density of the estimator. Using the estimated marginal likelihoods, we calculated the Bayes factor to assess model performance. Increases in the variance scaling increase the estimated Bayes factor, therefore we report a conservative Bayes factor bases on a small variance scaling S9.

### Comparing Codon Specific Parameter Estimates and Selecting Candidate lineages

As the choice of reference codon can reorganize codon families coding for an amino acid relative to each other, all parameter estimates were interpreted relative to the mean for each codon family.

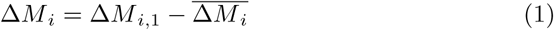

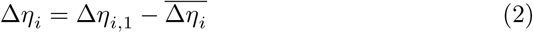

Comparison of codon specific parameters (Δ*M* and Δ*η* = 2*N*_*e*_q(*η*_*i*_ − *η*_*j*_)) was performed using the function lmodel2 in the R package lmodel2 (1.7.3) (Legendre, 2018) and R version 3.4.1 (R Core Team, 2013). The parameter Δ*η* can be interpreted as the difference in fitness between codon *i* and *j* for the average gene with *ϕ* = 1 scaled by the effective population size *N*_*e*_, and the selective cost of an ATP *q* (Gilchrist, 2007; Gilchrist *et al*., 2015). Type II regression was performed with re-centered parameter estimates, accounting for noise in dependent and independent variable (Sokal and Rohlf, 1981).

Due to the greater dissimilarity of the Δ*M* estimates between the endogenous and exogenous genes, and the slower decay rate of mutation bias, we decided to focus on our estimates of mutation bias to identify potential source lineages. The top ten lineages with the highest similarity in Δ*M* to the exogenous genes were selected as potential candidates (Figure 2).

### Phylogenetic Analysis

Using the dataset from Shen *et al*. (2018), we first identified 129 alignments for exogenous genes that further contained homologous genes for *E. gossypii*, and at least one other Lachancea species. We excluded all species from the alignments that do not belong to the Saccharomycetaceae clade. IQTree (Nguyen *et al*., 2015) was used to identify the best fitting model for each gene and to estimate the individual gene trees. Each gene tree was rooted using either *Saccharomyces cerevisiae, Saccharomyces uvarum, Saccharomyces eubayanus* as outgroup. We calculated the most recent common ancestor (MRCA) of *L. kluyveri* and *E. gossypii* as well as the MRCA of *L. kluyveri* and the remaining Lachancea. The distance between the MRCA and the root was used to asses which pairs (*L. kluyveri* and *E. gossypii*, or *L. kluyveri* and other Lachancea) have a more recent common ancestor.

### Synteny Comparison

We obtained complete genome sequences for all 10 candidate lineages (Table 2) from NCBI (on: 02-05-2017). Genomes were aligned and checked for synteny using SyMAP (4.2) with default settings (Soderlund *et al*., 2006, 2011). We assess synteny as percentage coverage of the exogenous gene region.

### Estimating Age of Introgression

We modeled the change in codon frequency over time using an exponential model for all two codon amino acids. While our approach is equivalent to Marais *et al*. (2004), we want to explicitly state the relationship between the change in codon frequency *c*_1_ as a function of mutation bias Δ*M* as

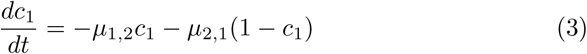

where *µ*_*i,j*_ is the rate at which codon *i* mutates to codon *j* and *c*_1_ is the frequency of the reference codon. Initial codon frequencies *c*_1_(0) for each codon family were taken from our mutation parameter estimates for *E. gossypii* where *c*_1_(0) = exp[Δ*M* _gos_]/(1 + exp[Δ*M* _gos_]). Our estimates of Δ*M* _endo_ can be used to calculate the steady state of equation 3 were 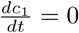to obtain the equality

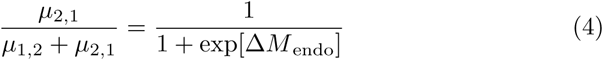

Solving for *µ*_1,2_ gives us *µ*_1,2_ = Δ*M* _endo_ exp[*µ*_2,1_] which allows us to rewrite and solve equation 3 as

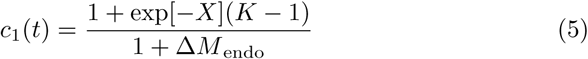

where *X* = (1 + Δ*M* _endo_)*µ*_2,1_*t* and *K* = *c*_1_(0)(1 + Δ*M* _endo_).

Equation 5 was solved with a mutation rate *µ*_2,1_ of 3.8 × 10^−10^ per nucleotide per generation (Lang and Murray, 2008). Current codon frequencies for each codon family where taken from our estimates of Δ*M* from the exogenous genes. Mathematica (11.3) (Wolfram Research Inc., 2017) was used to calculate the time *t*_intro_ it takes for the initial codon frequencies *c*_1_(0) for each codon family to equal the current exogenous codon frequencies. The same equation was used to determine the time *t*_decay_ at which the signal of the exogenous cellular environment has decayed to within 1% of the endogenous environment.

### Estimating Selection against Codon Mismatch

In order to estimate the selection against codon mismatch, we had to make three key assumptions. First, we assumed that the current exogenous amino acid sequence of a gene is representative of its ancestral state and the replaced endogenous gene it replaced. Second, we assume that the currently observed cellular environment of *E. gossypii* reflects the cellular environment that the exogenous genes experienced before transfer to *L. kluyveri*. Lastly, we assume that the difference in the efficacy of selection between the cellular environments due to differences in either effective population size *N*_*e*_ or the selective cost of an ATP *q* of the source lineage and *L. kluyveri* can be expressed as a scaling constant and that protein synthesis rate *ϕ* has not changed between the replaced endogenous and the introgressed exogenous genes. Using estimates for *N*_*e*_ = 1.36 × 10^7^ (Wagner, 2005) for *Saccharomyces paradoxus* we scale our estimates of Δ*η* which explicitly contains the effective population size *N*_*e*_ (Gilchrist *et al*., 2015) and define 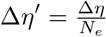.

All of our genome parameter estimations are scaled by lineage specific effects such as *N*_*e*_, the average, absolute gene expression level, and/or the proportionate fitness value of an ATP. In order to account for these genome specific differences in scaling, we scale the difference in the efficacy of selection on codon usage between the donor lineage and *L. kluyveri* using a linear scaling factor *κ*. As Δ*η* is defined as Δ*η* = 2*N*_*e*_q(*η*_*i*_ − *η*_*j*_), we cannot distinguish if *κ* is a scaling on protein synthesis rate *ϕ*, effective population size *N*_*e*_, or the selective cost of an ATP *q* (Gilchrist, 2007; Gilchrist *et al*., 2015). We calculated the selection against each genes codon mismatch assuming additive fitness effects as

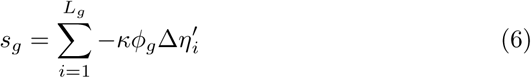

where *s*_*g*_ is the overall strength of selection for translational efficiency on gene, *g* in the exogenous gene set, *κ* is a constant, scaling the efficacy of selection between the endogenous and exogenous cellular environments, *L*_*g*_ is length of the protein in codons, *ϕ*_*g*_ is the estimated protein synthesis rate of the gene in the endogenous environment, and 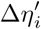, is the Δ*η*′ for the codon at position *i*. As stated previously, our Δ*η* are relative to the mean of the codon family. We find that the selection against the introgressed genes is minimized at *κ* ∼ 5 (Figure S7b). Thus, we expect a five fold difference in the efficacy of selection between *L. kluyveri* and *E. gossypii*, due to differences in either protein synthesis rate *ϕ*, effective population size *N*_*e*_, and/or the selective cost of an ATP *q*. Therefore, we set *κ* = 1 if we calculate the *s*_*g*_ for the endogenous and the current exogenous genes, and *κ* = 5 for *s*_*g*_ for selection calculations at the time of introgression.

However, since we are unable to observe codon sequences of the replaced endogenous genes and for the exogenous genes at the time of introgression, instead of summing over the sequence, we calculate the expected codon count *E*[*n*_*g,i*_] for codon *i* in gene *g* simply as the probability of observing codon *i* multiplied by the number of times the corresponding amino acids is observed in gene *g*, yielding:

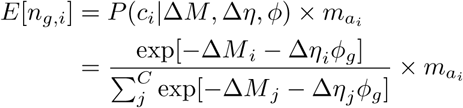

where 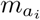 is the number of occurrences of amino acid *a* that codon *i* codes for. Thus replacing the summation over the sequence length *L*_*g*_ in equ. (6) by a summation over the codon set *C* and calculating *s*_*g*_ as

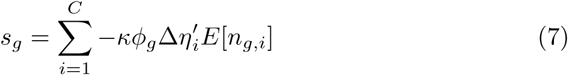

We report the selection due to mismatched codon usage of the introgression as Δ*s*_*g*_ = *s*_intro,*g*_ − *s*_endo,*g*_ where *s*_intro,*g*_ is the selection against an introgressed gene *g* either at the time of the introgression or presently.

### Randomizing genes

We randomized the codon content of the endogenous and exogenous genes while conserving the di-nucleotide distribution and GC content using the randomization algorithm from SPARCS (Zhang *et al*., 2013). We used the default settings of the randomization algorithm. The resulting gene sets were analyzed using the same scheme as described above.

## Availability of data and materials

Parameter estimates generated during this study are available from the corresponding author. All remaining data generated during this study are included in this published article as figures, and tables.

## Authors’ contributions

CL and MAG initiated the study. CL collected and analyzed the data and wrote the manuscript. MAG and BCO edited the manuscript. CL, MAG, BCO, and RZ contributed to the data analysis and acquiring of funding. All Authors approved the final manuscript.

## Acknowledgments

The authors would like to thank Alexander Cope for helpful criticisms and suggestions for this work.

## Supplementary Material

Supporting Materials for *Unlocking a signal of introgression from codons in Lachancea kluveri using a mutation-selection model by Landerer et al..*

**Table S1:** Synonymous mutation codon preference based on our estimates of Δ*M*. Shown are the most likely codon in low expression genes for each amino acid in: *E. gossypii*, in the endogenous and exogenous genes of *L. kluyveri*, and in the combined *L. kluyveri* genome without accounting for the two cellular environments.

| Amino Acid | *E. gossypii* | Endogenous | Exogenous | Combined |
| --- | --- | --- | --- | --- |
| Ala A | GCG | GCA | GCG | GCG |
| Cys C | TGC | TGT | TGC | TGC |
| Asp D | GAC | GAT | GAC | GAC |
| Glu E | GAG | GAA | GAG | GAG |
| Phe F | TTC | TTT | TTT | TTT |
| Gly G | GGC | GGT | GGC | GGC |
| His H | CAC | CAT | CAC | CAC |
| Ile I | ATC | ATT | ATC | ATA |
| Lys K | AAG | AAA | AAG | AAA |
| Leu L | CTG | TTG | CTG | CTG |
| Asn N | AAC | AAT | AAC | AAT |
| Pro P | CCG | CCA | CCG | CCG |
| Gln Q | CAG | CAA | CAG | CAG |
| Arg R | CGC | AGA | AGG | CGG |
| Ser<sub>4</sub> S | TCG | TCT | TCG | TCG |
| Thr T | ACG | ACA | ACG | ACG |
| Val V | GTG | GTT | GTG | GTG |
| Tyr Y | TAC | TAT | TAC | TAC |
| Ser<sub>2</sub> Z | AGC | AGT | AGC | AGC |

**Table S2:** Synonymous selection codon preference based on our estimates of Δ*η*. Shown are the most likely codon in high expression genes for each amino acid in: *E. gossypii*, in the endogenous and exogenous genes of *L. kluyveri*, and in the combined *L. kluyveri* genome without accounting for the two cellular environments.

| Amino Acid | *E. gossypii* | Endogenous | Exogenous | Combined |
| --- | --- | --- | --- | --- |
| Ala A | GCT | GCT | GCT | GCT |
| Cys C | TGT | TGT | TGT | TGT |
| Asp D | GAT | GAC | GAT | GAT |
| Glu E | GAA | GAA | GAA | GAA |
| Phe F | TTT | TTC | TTC | TTC |
| Gly G | GGA | GGT | GGT | GGT |
| His H | CAT | CAC | CAT | CAT |
| Ile I | ATA | ATC | ATT | ATT |
| Lys K | AAA | AAG | AAA | AAG |
| Leu L | TTA | TTG | TTG | TTG |
| Asn N | AAT | AAC | AAT | AAC |
| Pro P | CCA | CCA | CCT | CCA |
| Gln Q | CAA | CAA | CAA | CAA |
| Arg R | AGA | AGA | AGA | AGA |
| Ser<sub>4</sub> S | TCA | TCC | TCT | TCT |
| Thr T | ACT | ACC | ACT | ACT |
| Val V | GTT | GTC | GTT | GTT |
| Tyr Y | TAT | TAC | TAT | TAC |
| Ser<sub>2</sub> Z | AGT | AGT | AGT | AGT |

**Figure S1:**
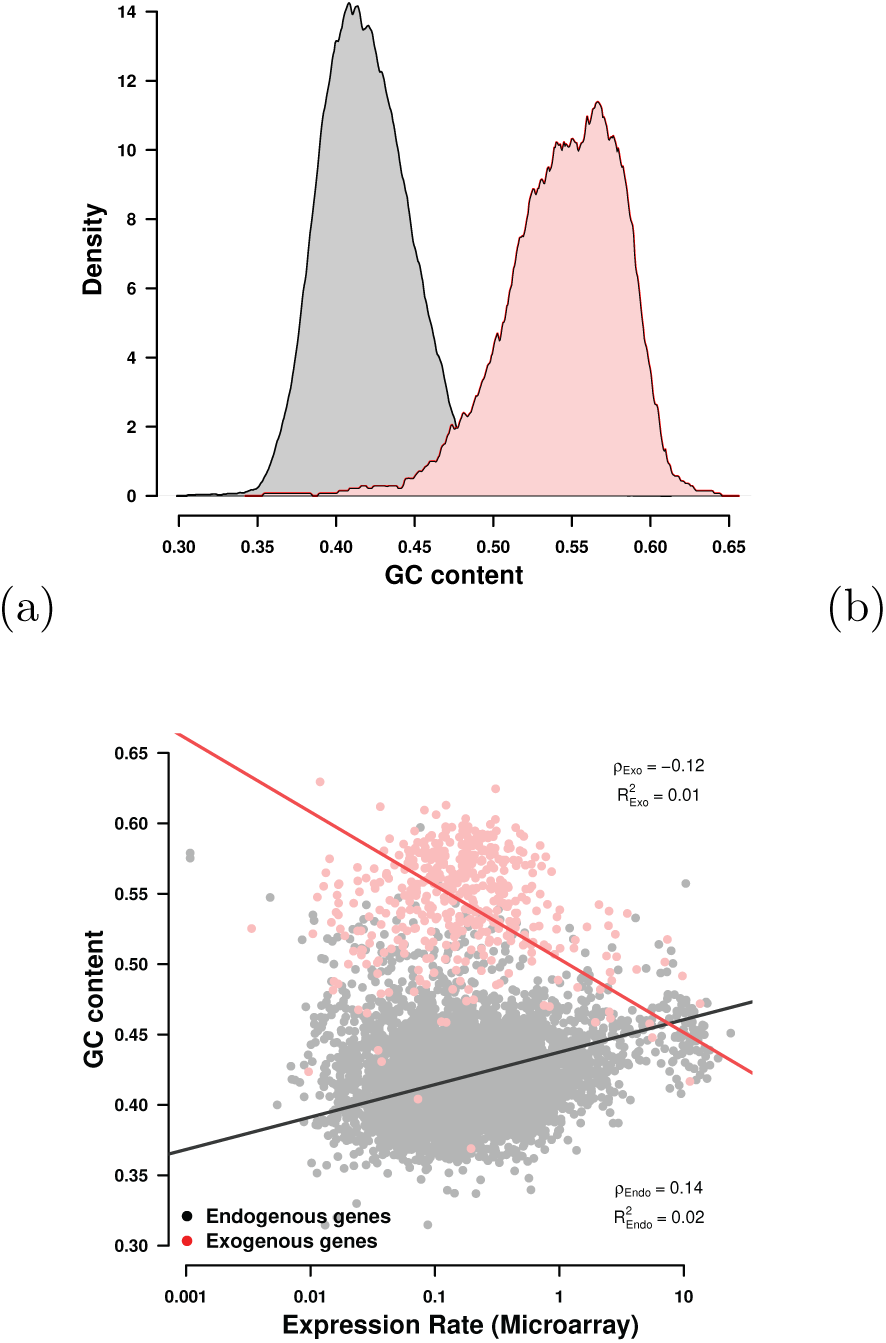
Endogenous and exogenouns genes have distinct GC content. (a) Distribution of GC content content in the endogenous and exogenous genes. (b) Correlation of endogenous and exogenous GC content with measured gene expression. While the endogenous GC content shows a slight positive correlation with gene expression (*ρ* = 0.14, *p* = 1.2 × 10^−21^), the exogenous GC content is negatively correlated with gene expression (*ρ* = −0.12, *p* = 0.014).

**Figure S2:**
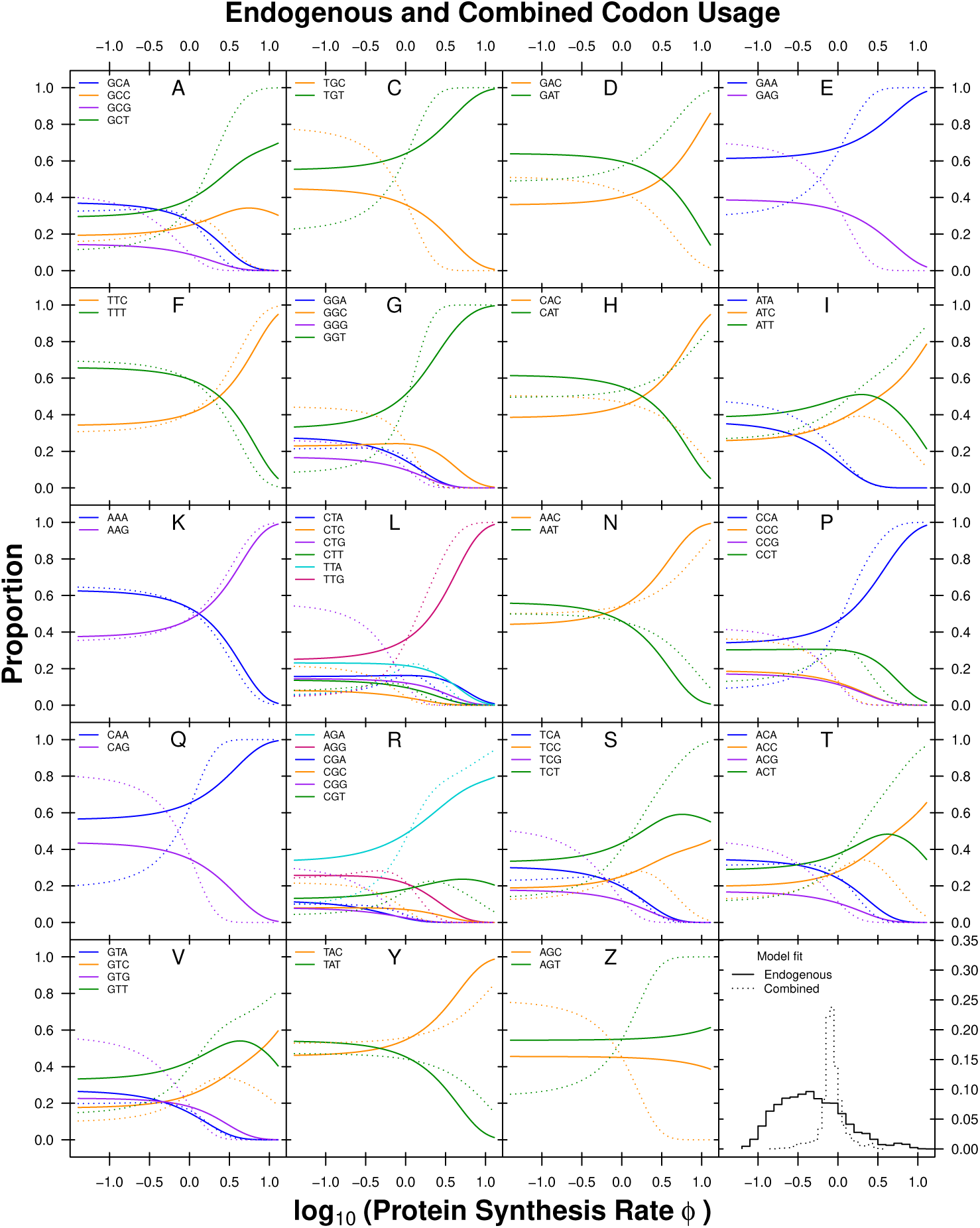
Codon usage patterns for 19 amino acids. Amino acids are indicated as one letter code. The amino acids Serine was split into two groups (S and Z) as Serine is coded for by two groups of codons that are separated by more than one mutation. Solid line indicates the endogenous codon usage, dotted line indicates the combined codon usage.

**Figure S3:**
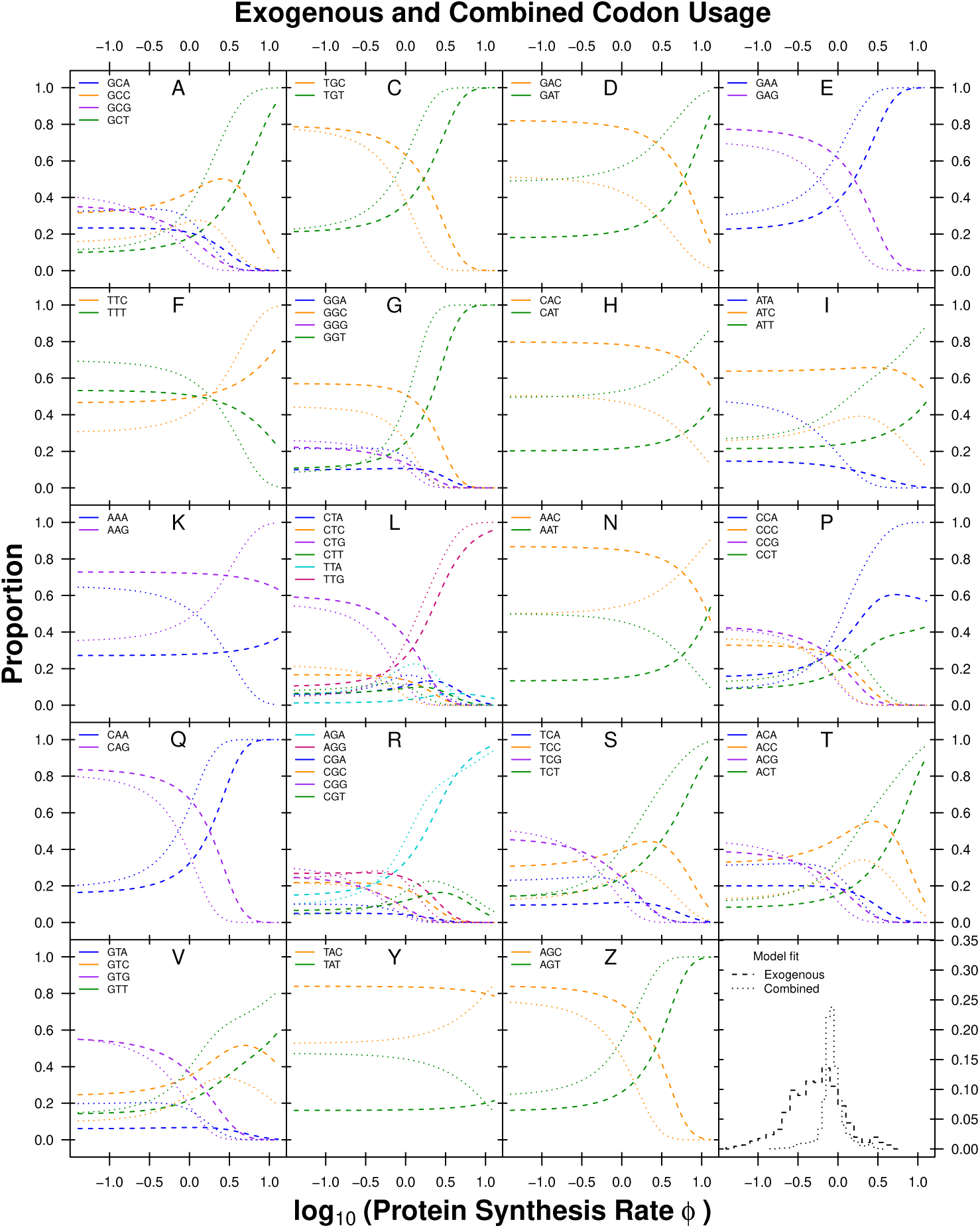
Codon usage patterns for 19 amino acids. Amino acids are indicated as one letter code. The amino acids Serine was split into two groups (S and Z) as Serine is coded for by two groups of codons that are separated by more than one mutation. dashed line indicates the exogenous codon usage, dotted line indicates the combined codon usage.

**Figure S4:**
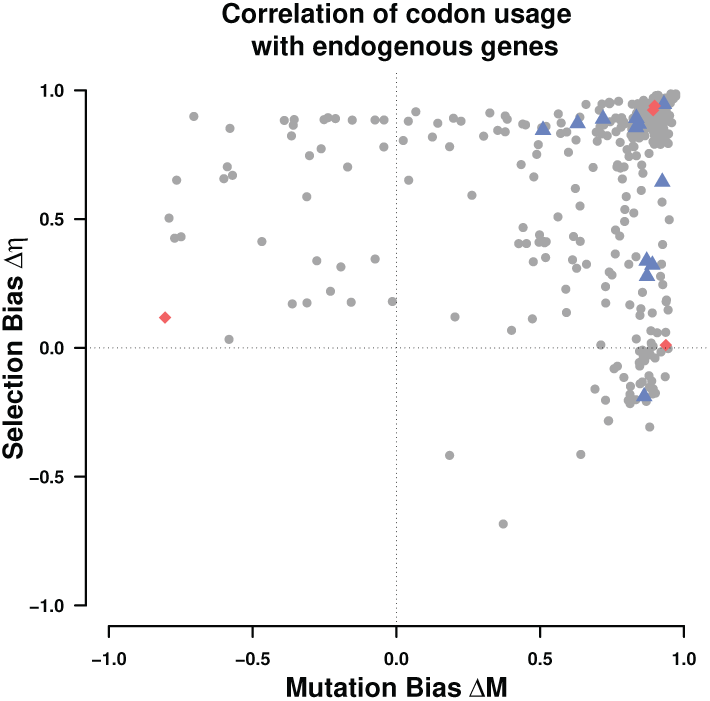
Correlation coefficients of Δ*M* and Δ*η* of the endogenous genes with 332 examined budding yeast lineages. Dots indicate the correlation of Δ*M* and Δ*η* of the lineages with the exogenous parameter estimates. Blue triangles indicate the Lachancea and red diamonds indicate Eremothecium lineages. All regressions were performed using a type II regression (Sokal and Rohlf, 1981).

**Figure S5:**
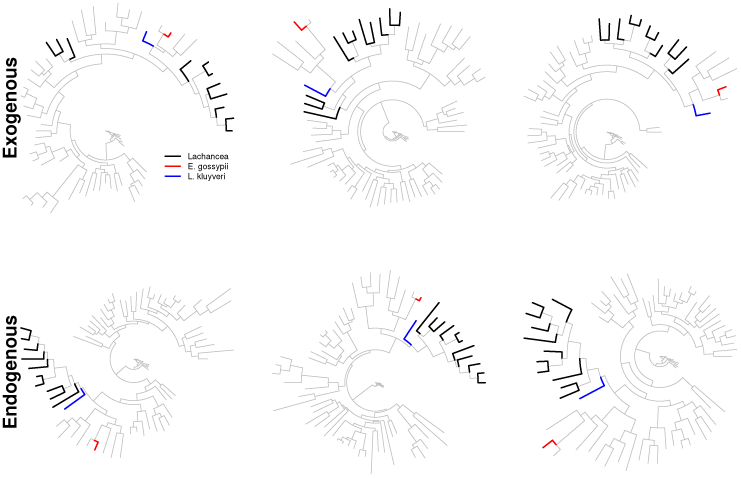
Gene trees illustrating the placement of *L. kluyveri* (blue) and *E. gossypii* (red) for three endogenous and three exogenous genes. The remaining Lachancea are highlighted in black. (Top row) Gene trees for three exogenous genes (from left to right: SAKL0C05742g, SAKL0C03520g, SAKL0C02376g). (Bottom row) Gene trees for three endogenous genes (from left to right: SAKL0D03960g, SAKL0G02354g, SAKL0H02552g).

**Figure S6:**
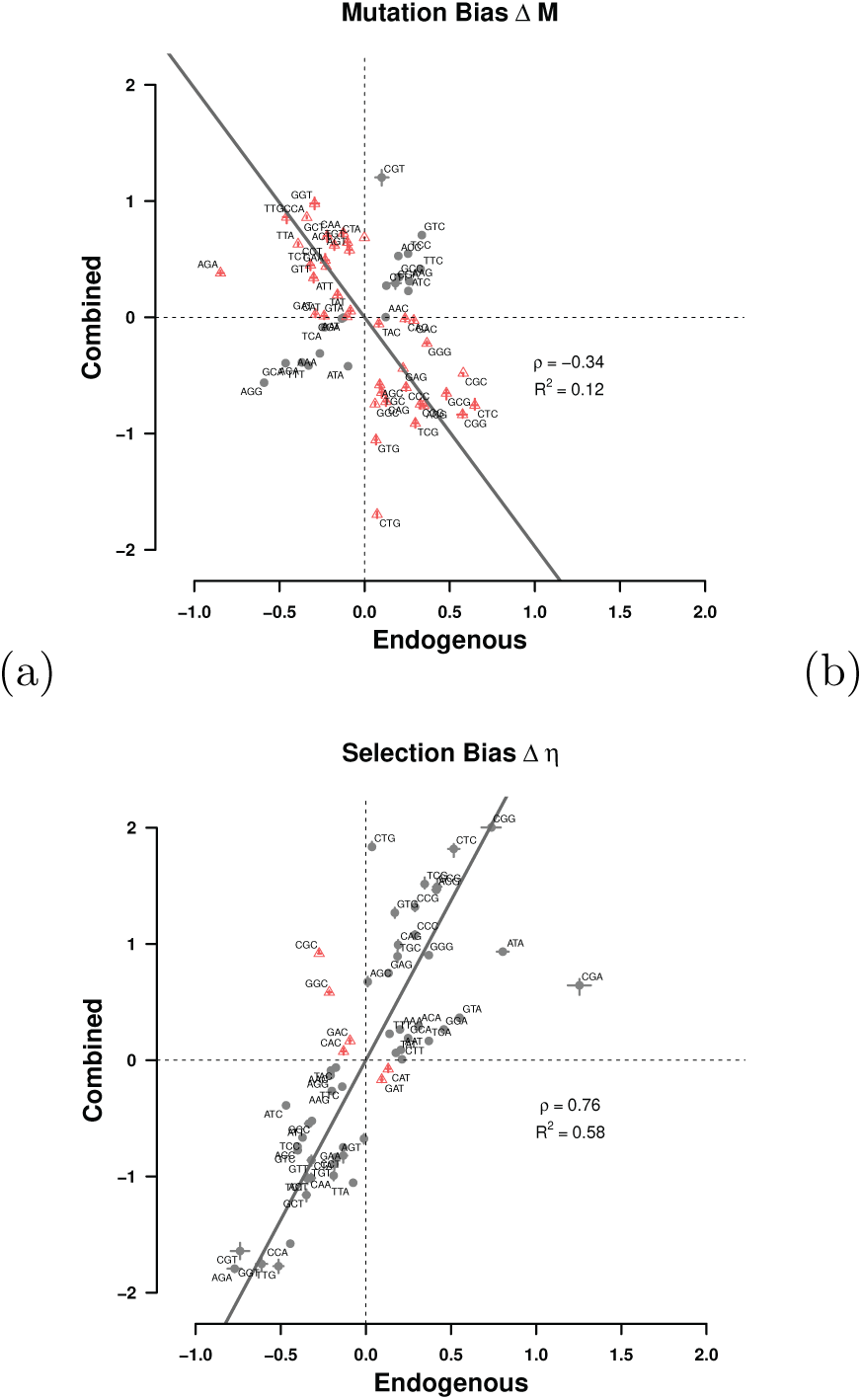
Comparison of (a) mutation bias Δ*M* and (b) selection bias Δ*η* parameters for endogenous genes and combined gene sets. Estimates are relative to the mean for each codon family. Black dots indicate Δ*M* or Δ*η* parameters with the same sign for the endogenous and exogenous genes, red dots indicate parameters with different signs. Black line indicates type II regression line (Sokal and Rohlf, 1981). Dashed lines mark quadrants.

**Figure S7:**
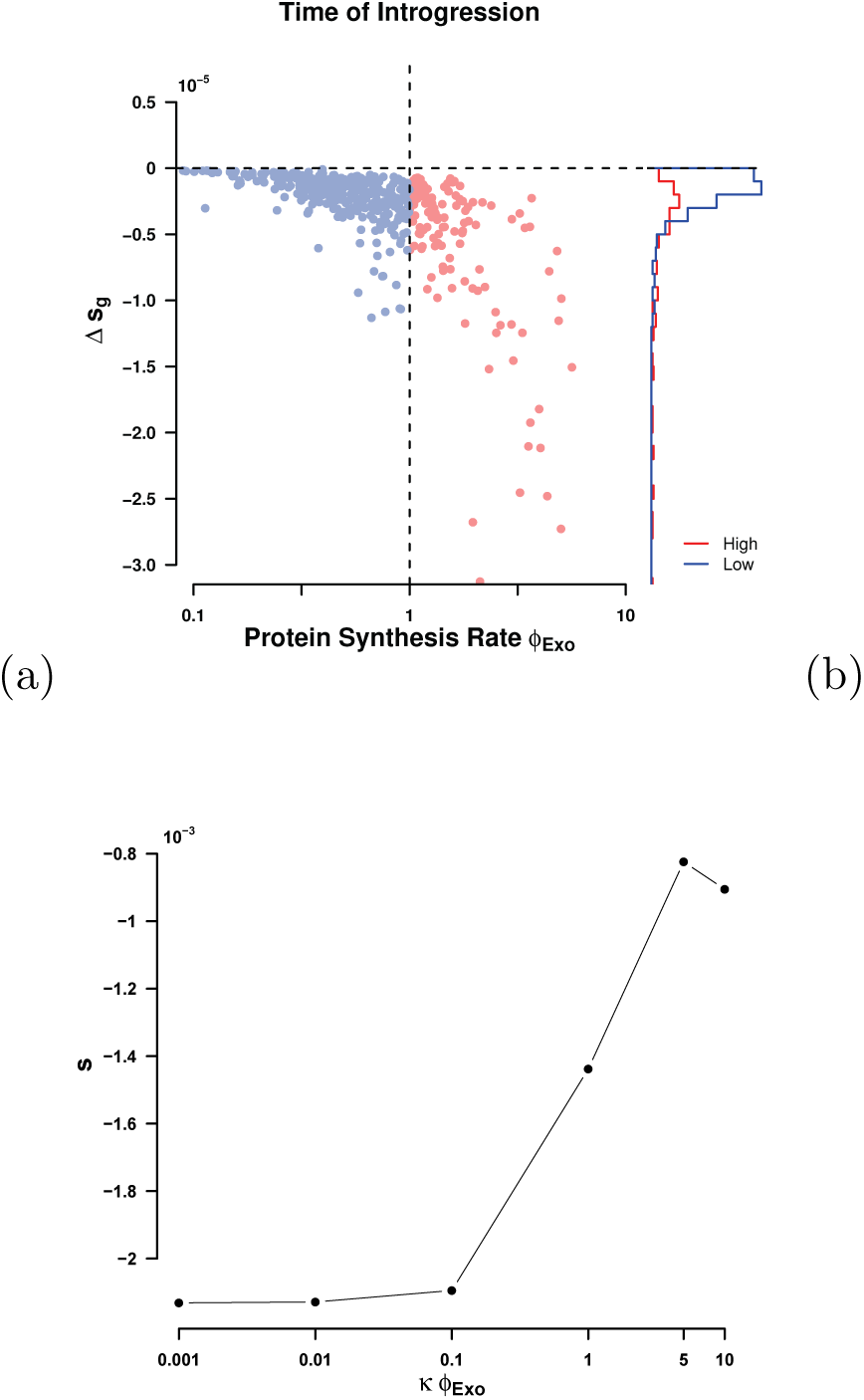
Selection against mismatched codon usage (a) without scaling of *ϕ* per gene. Vertical dashed line indicates split between high and low expression genes at *ϕ* = 1. Horizontal dashed line indicates neutrality. (b) Change of total selection against mismatched codon usage with scaling term *κ* between *E. gossypii* and *L. kluyveri*

**Figure S8:**
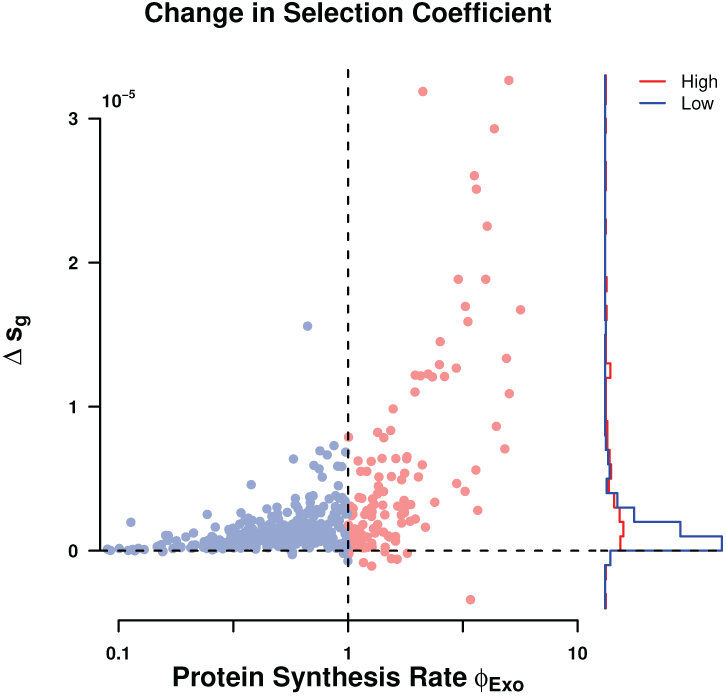
Total amount of adaptation estimated to have occurred between time of introgression and currently observed per gene. Vertical dashed line indicates split between high and low expression genes at *ϕ* = 1. Horizontal dashed line indicates no change in selection against mismatched codon usage.

**Figure S9:**
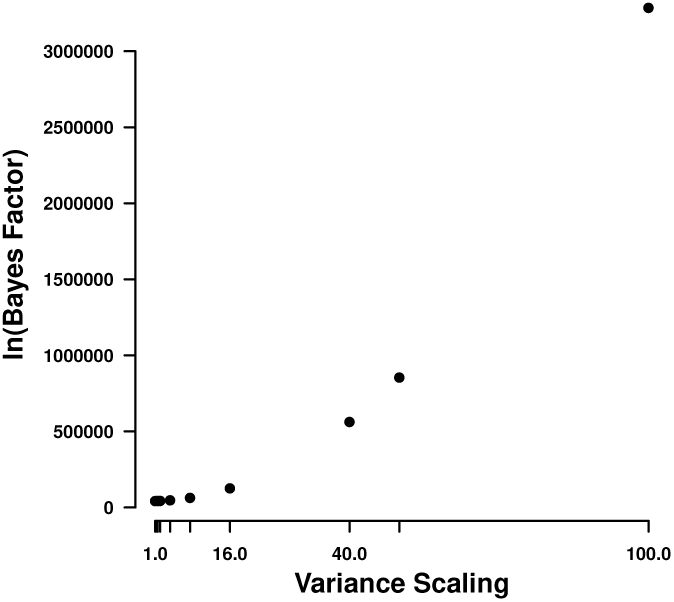
Influence of the variance scaling of the importance distribution on the estimated Bayes factor.

**Figure S10:**
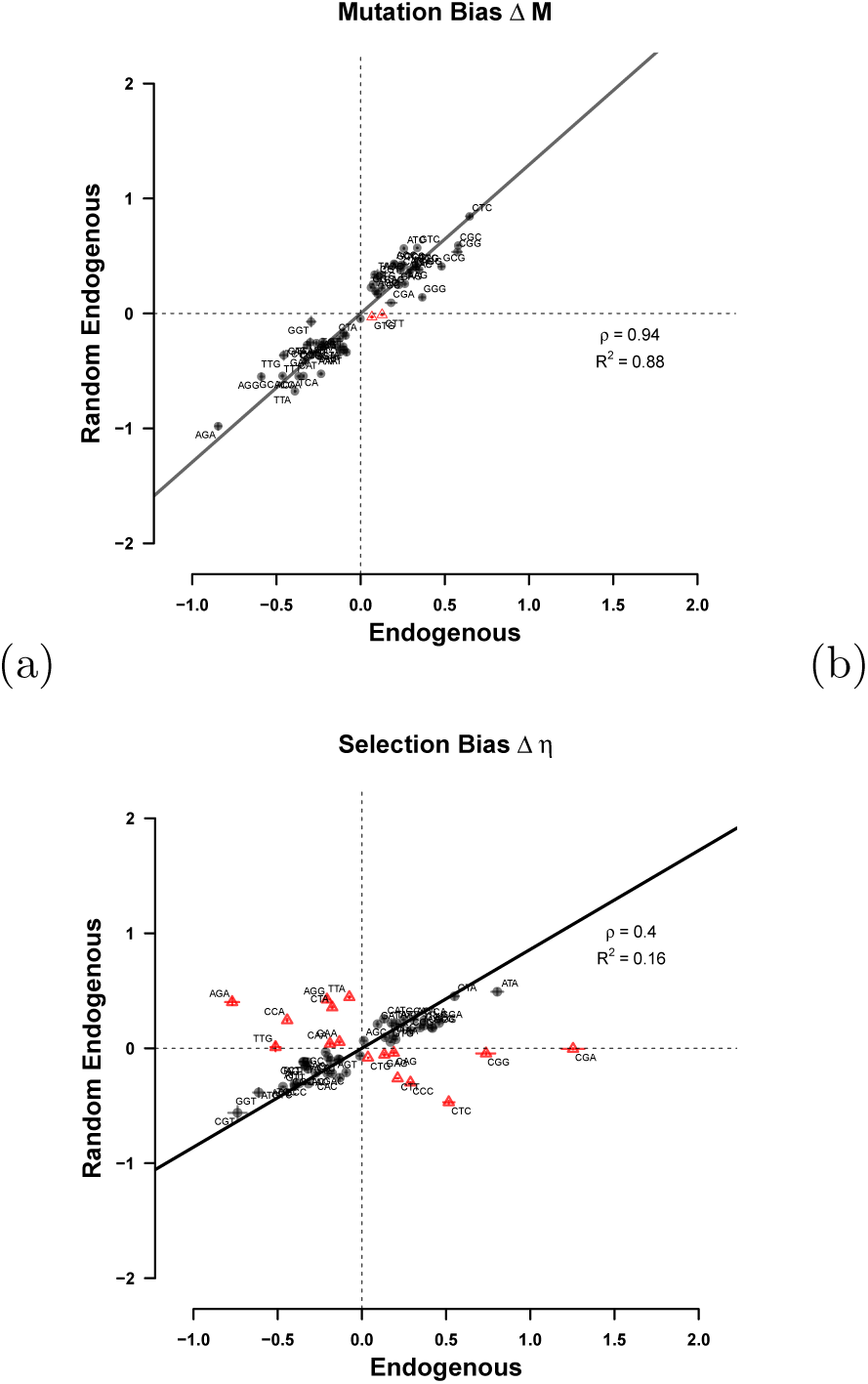
Comparison of mutation bias Δ*M* (a) and selection bias Δ*η* (b) estimated from the endogenous genes and their randomized counterparts.

**Figure S11:**
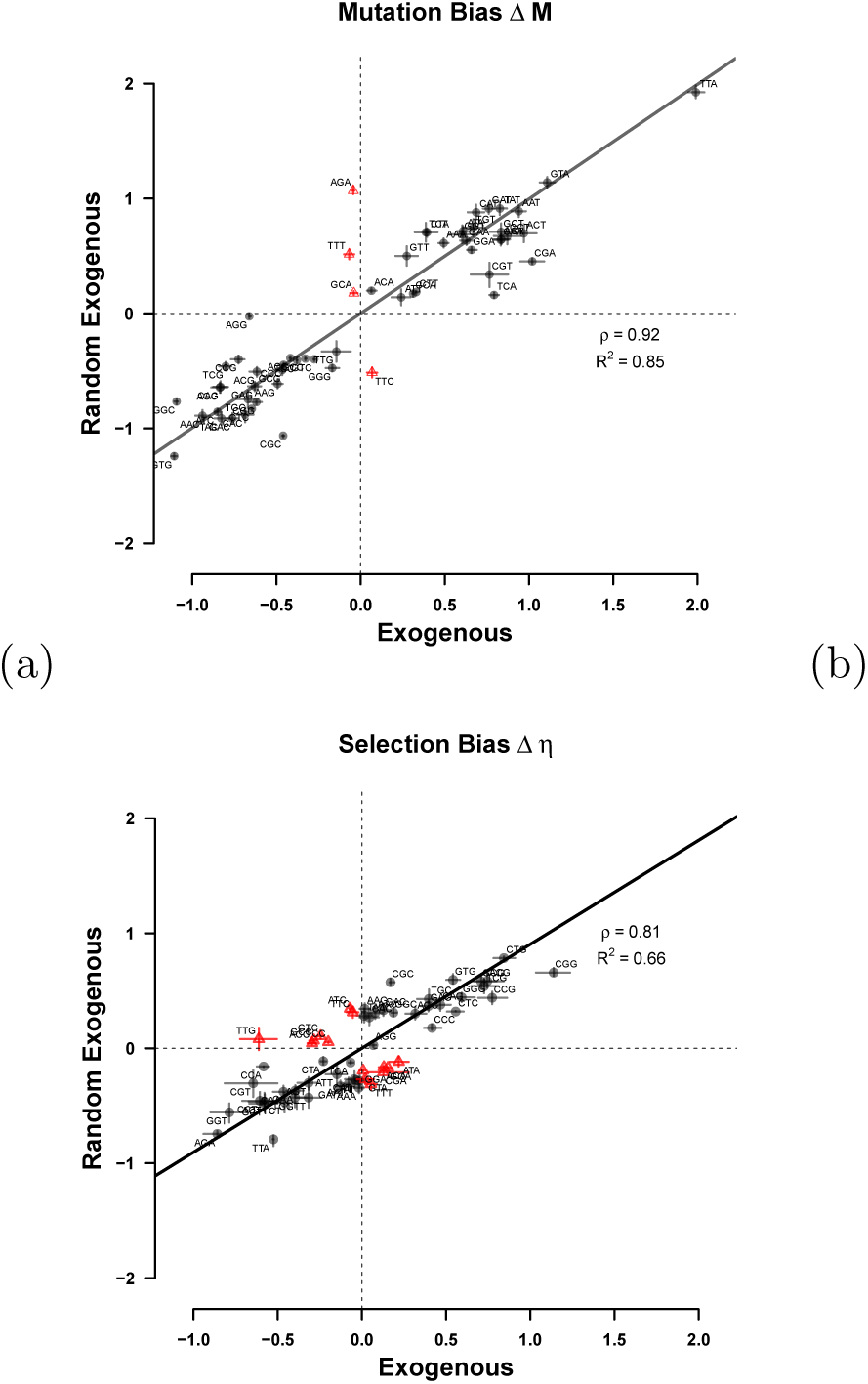
Comparison of mutation bias Δ*M* (a) and selection bias Δ*η* (b) estimated from the exogenous genes and their randomized counterparts.

**Figure S12:**
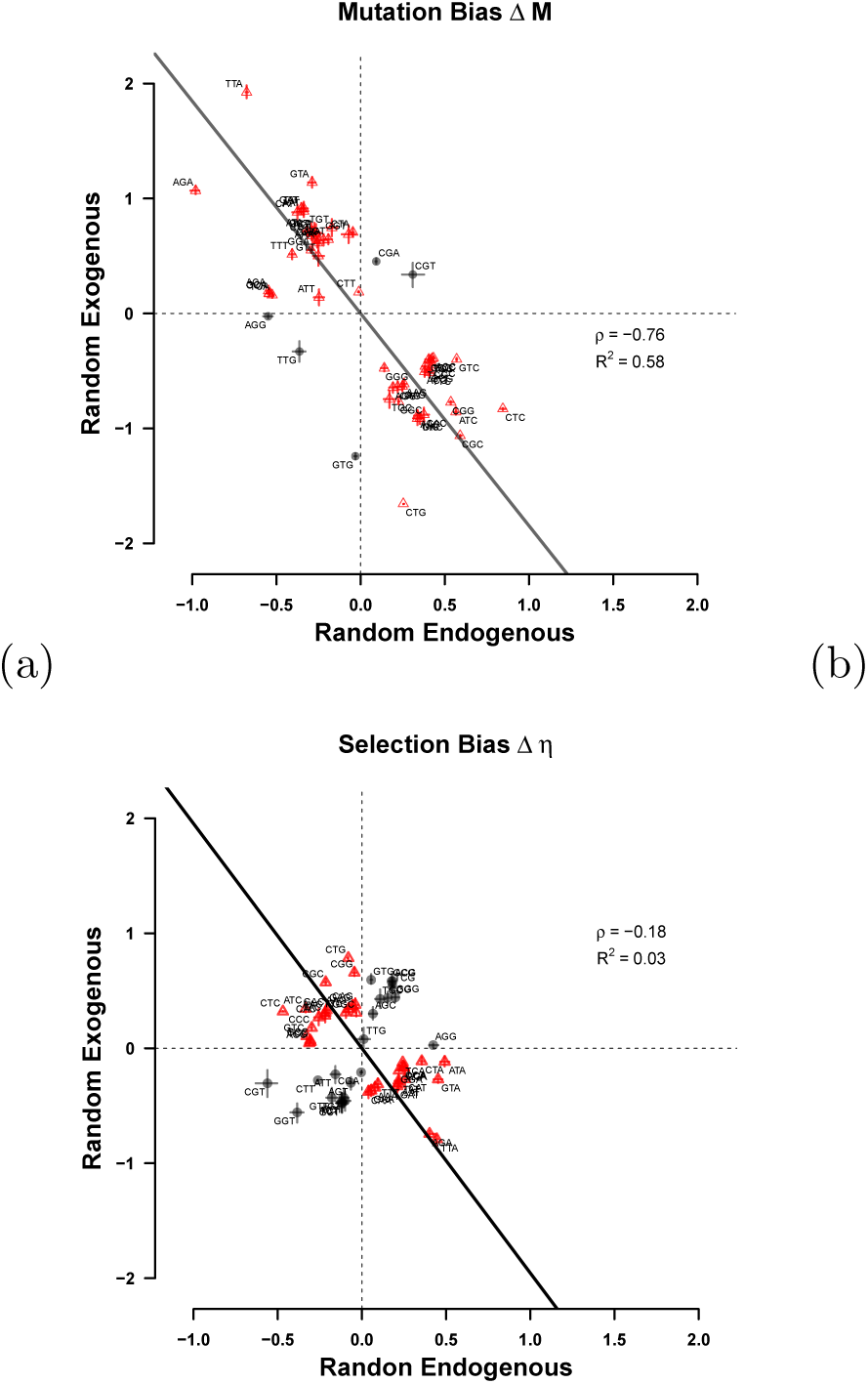
Comparison of mutation bias Δ*M* (a) and selection bias Δ*η* (b) estimated from the randomized endogenous and the di-nucleotide randomized exogenous genes.

**Figure S13:**
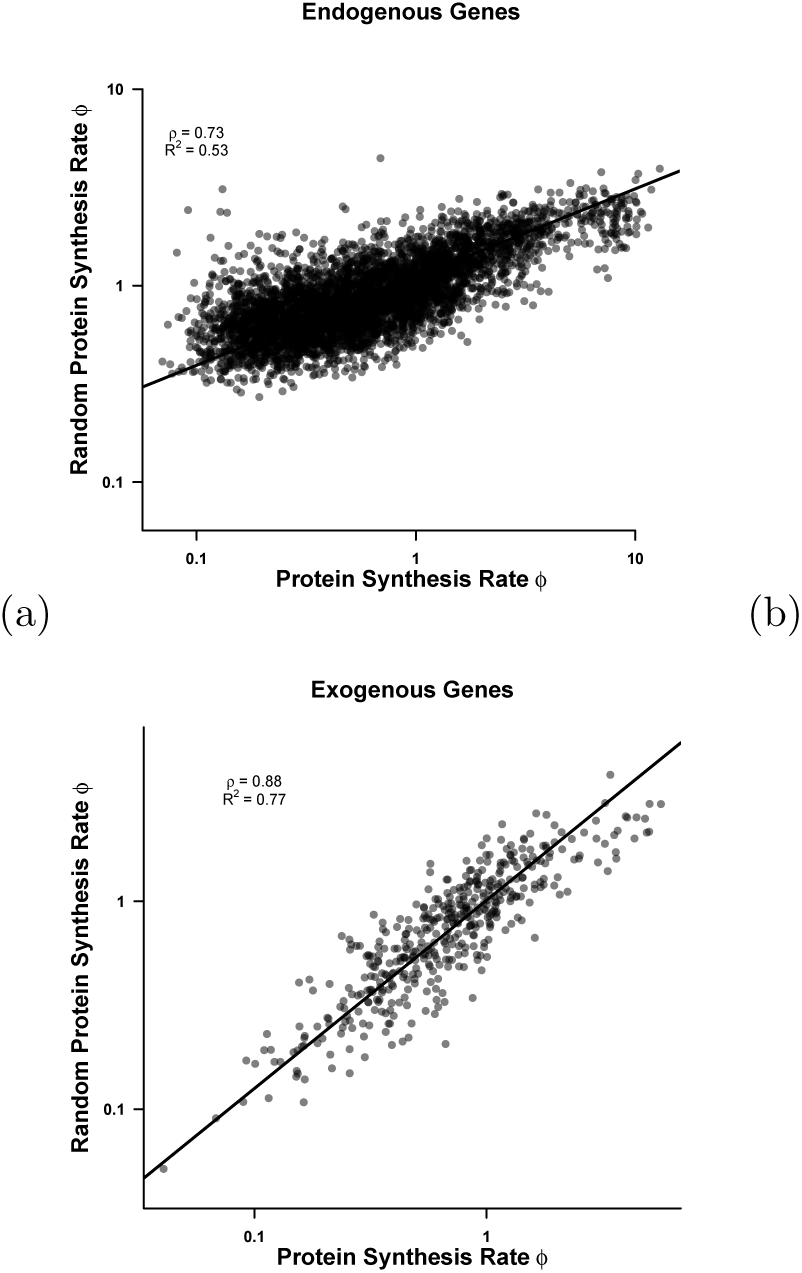
Comparison of protein synthesis rate *ϕ* estimated from the endogenous (a) and exogenous genes (b) and their randomized counterparts.

